# Gossamer: Scaling Image Processing and Reconstruction to Whole Brains

**DOI:** 10.1101/2024.04.07.588466

**Authors:** Karl Marrett, Keivan Moradi, Chris Sin Park, Ming Yan, Chris Choi, Muye Zhu, Masood Akram, Sumit Nanda, Qing Xue, Hyun-Seung Mun, Adriana E. Gutierrez, Mitchell Rudd, Brian Zingg, Gabrielle Magat, Kathleen Wijaya, Hongwei Dong, X. William Yang, Jason Cong

## Abstract

Neuronal reconstruction–a process that transforms image volumes into 3D geometries and skeletons of cells– bottlenecks the study of brain function, connectomics and pathology. Domain scientists need *exact* and *complete* segmentations to study subtle topological differences. Existing methods are disk-bound, dense-access, coupled, single-threaded, algorithmically unscalable and require manual cropping of small windows and proofreading of skeletons due to low topological accuracy. Designing a data-intensive parallel solution suited to a neurons’ shape, topology and far-ranging connectivity is particularly challenging due to I/O and load-balance, yet by abstracting these vision tasks into strategically ordered specializations of search, we progressively lower memory by 4 orders of magnitude. This enables 1 mouse brain to be fully processed in-memory on a single server, at 67× the scale with 870× less memory while having 78% higher automated yield than APP2, the previous state of the art in performant reconstruction.

## I. Introduction

Lightsheet microscopy [43] has reached a maturity that it can digitize whole mouse brains in 1 day and produce terabytes of data that require image processing and reconstruction (Figure 1). Unfortunately, these new datasets, at baseline, have marginal value since they lack quality and semantic labeling. How do we future-proof software design against the increasing scale of unstructured data to derive meaningful 3D assets? We offer an application that can efficiently transform large-scale volumes into relatively accurate models for analysis.

**Fig. 1.**
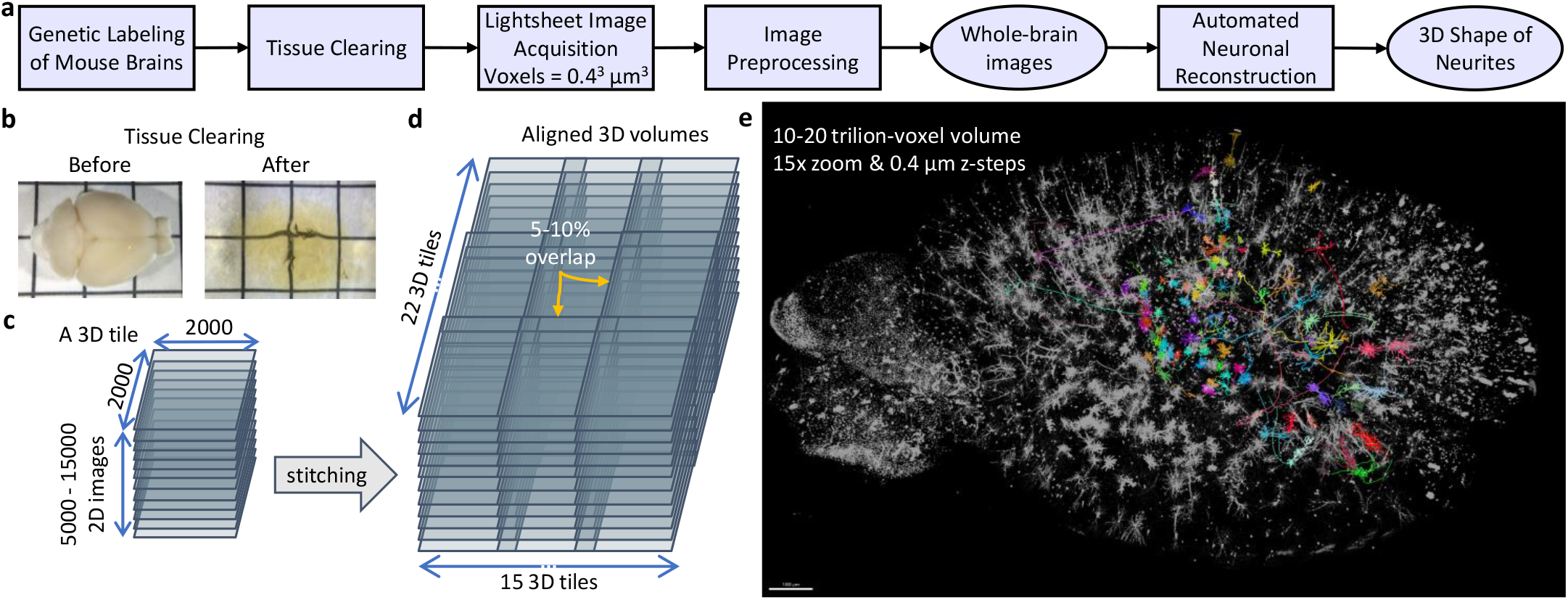
Whole-brain lightsheet data. (a) Data acquisition and processing pipeline. (b) A sample mouse brain before and after tissue clearing. (c) Since the field of view of the microscope is limited and cannot cover the entire brain in one scan lightsheet data is in the form of a 2D image series that form 3D tiles. (d) Image stitching is needed to make a whole-brain image. (e) A 3D rendering of the image (black and white) and some of the reconstructed neurons (colored part) that shows the scale of the problem.

Intelligent systems (e.g., biological brains, computer microprocessors) have inherent long-range context that benefits from reconstructing as a complete whole. However, existing methodologies can not scale to modern data sizes; we test up to 30 TB raw inputs for this study. Unstructured data has a larger memory footprint and necessitates further computation to extract a useful representation, yet data bandwidth is a bottleneck across all parts of the modern computing infrastructure stack. This makes distributed or cloud strategies impractical, suggesting instead an edge computing approach. Therefore, our design proposes a single system composed of 1 microscope and 1 workstation processing a single brain at a time. This is feasible because a single modern server can have terabytes of memory and multiple CPUs and GPUs with hundreds of cores. Our system can efficiently process or reconstruct a whole brain in a similar time that it takes to image or transfer to the cloud (about 4 days each).

Lack of large labeled datasets of 3D geometries is also an issue in the broader graphics community but this need is more acute in science. For researchers to take advantage of recent data-driven and supervised methods they need highly specific, often novel, training labels tailored to new scientific queries. A further complication is that, unlike in other domains like the arts, accuracy is paramount. Neuroscientists will go to great lengths to ensure that acquired data reflects ground truth exactly before using it for scientific analysis since experimental differences can be incredibly subtle. Transforming automated but inaccurate labels into fully correct ones, which we refer to as *proofreading*, is a time-consuming interactive human-led process. Understandably, this creates a bottleneck where acquiring the numerous proofread samples required for statistical significance prolongs project iteration time.

Brain research has a constricting limitation: reconstruction. This involves (1) image processing and encoding (2) segmenting cell bodies (seeds) (3) segmenting long-range branches and (4) transforming into *curve skeletons* [18]. Morphology, the 3D shape of semantic objects like neurons, and topology, the arrangement, coverage and connectivity, may be a contributing facet in the study of function, pathology and cell type [51]. *Reconstruction*, a common method of studying neuronal morphology, is a process in which fluorescently labeled cells in 3D images are segmented and compacted into ball and stick models (skeletons) [14]. Skeletons are uniquely useful in shape analysis as they distill both topological and morphological features. Recent reviews in neuroscience identified that a scalable [3], comprehensive application [30] for single-cell reconstruction is the greatest need of the community.

Gossamer is an end-to-end pipeline for image processing and reconstruction of lightsheet images with several unique advancements.

1. We introduce several novel 3D and adaptive methods and accomplish enhancement, artifact removal, segmentation and reconstruction of high-scale images.
2. We achieve state of the art (SOTA) performance in space and time even when compared to the equivalent 2D SOTA via linear complexity algorithms, which is critical for rhizomatic structures (neurons) that are power-law in size.
3. We advance spatially sparse methodologies to enable the long-range contexts inherent in neuroscience to be computed entirely in-memory on a single workstation.
4. Across many of our workloads, we have the first hardware-accelerated and parallel implementations for multi-GPU or multi-core environments.
5. We enable efficient scaling to whole brain volumes (13000×13000×16000 voxels), a 67× end-to-end size increase over the most scalable alternate algorithm.

## II. Background and Related Work

Alternative methods fall short in these critical areas which leads to severe losses in automation, yield, performance and accuracy.

### A. Lightsheet imaging data

Raw lightsheet data consists of 3D tiles that must be preprocessed to remove camera artifacts, stitched together to form a whole volume, and post-processed to normalize pixel intensity (brightness) and refine the resolution. There are three major sources of non-uniformity in image brightness [39]: (1) some CMOS cameras have midline artifacts that produce a vertical artifact in the center of each tile. On the horizontal axis, there are also horizontal non-uniformities in brightness; (2) some objects in the tissue can partially block the passage of laser light creating horizontal shadows, i.e. *stripes*; (3) repeated imaging of areas desaturates the brightness unevenly referred to as *image bleaching*.

It is better to address CMOS artifacts, shadows and inherent image noise before stitching the image tiles to form a whole volume since these issues are confined to each tile, the computational cost is higher after stitching, and correcting them can improve automated alignment of tiles. However, correcting image bleaching after stitching is preferred since it benefits from a wider view of the whole brain image. Lightsheet, like other light microscopy forms, suffers from a lower resolution and a blur of signal on the z-axis. Deconvolution not only mitigates this issue but also increases foreground signal brightness. By addressing global image normalization and enhancing the foreground signal, a simple thresholding algorithm can segment the image (Figure 5a). Although existing software can function at smaller scales [44][11][9], we are not aware of a complete software pipeline that can operate at the native scale of modern lightsheet microscopy.

### B. Seed segmentation

Segmentation spatially assigns contiguous pixels of an image that belong to classes with a known meaning, for example, cell bodies. These filled regions can also be broken into instances, separated by occurrence in the volume. Neural network models that leverage convolution like U-net [40] are adept at these tasks at small scales. They also are linear time complexity but have poor runtime and high resource usage due to their space complexity which is on the order of the whole image–i.e., the working set W–which is prohibitive.

### C. Cell segmentation

In addition to some unavoidable level of image artifacts, neurons branches are thin, intricate and filament-like. This poses a challenge in automated segmentation and topological accuracy. If the frequency or severity of errors is high, humans will simply discard a neuron or start reconstructing from scratch rather than invest the time to proofread an automated output. We refer to the rate of adequately accurate outputs as the *automated yield*.

Though the problems in neuron reconstruction are specific, the metrics used do not adequately differentiate the causes of errors. We categorize relevant issues into semantics, morphology and topology as shown in Figure 2. *Semantics* relates to errors in the detection or classification of domain-specific structures. *Morphology* refers to errors in surface reconstruction such as breaks, collisions or other unfaithful aspects of the segmented region. *Topology* describes errors in the automatically reconstructed tree graphs when compared to their ground truth skeletons. Curve skeletons reveal both morphological and topological details of objects.

**Fig. 2.**
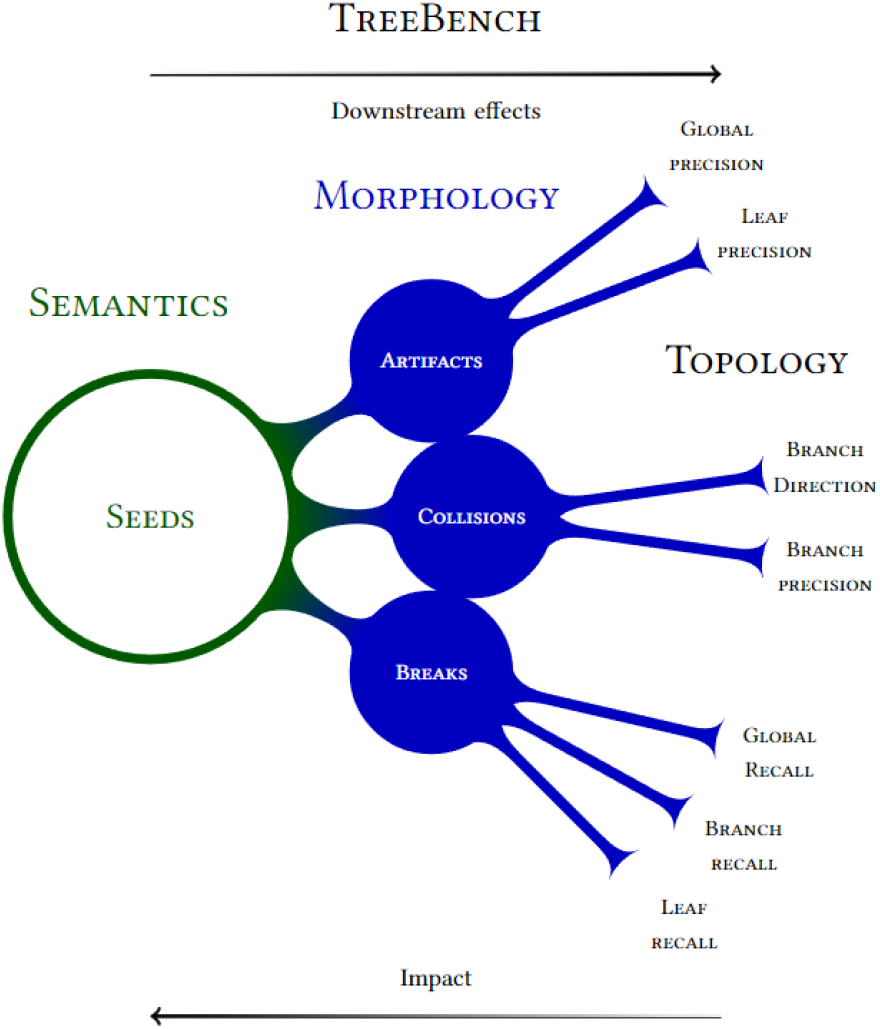
A schematic of various error types and their relative importance. The diagram itself loosely resembles the topological structure of a cell.

In this work, there are only 2 biological semantic categories: cell bodies (shown in green) and branches (blue) as illustrated in Figure 2. Cell bodies, also known as *somas*, are the root of a neuron’s branching projections. Due to their importance in neuronal reconstruction, it is clear that somas are natural *seeds* (the starting set) in the search for all connected branches. Semantic classification and morphological segmentation can be evaluated with established metrics such as recall and precision which report rates of false negatives and false positives respectively.

Tree topological errors are more nebulous and usually have an upstream cause from the segmentation input to skeletonization. *Path breaks* are false discontinuities in the segmented branches of cells. In contrast, over-segmentation leads to *path collisions*–the incorrect joining of disconnected branches or neuronal paths, which are biologically infeasible. Collisions can either be *intra* (within) or *inter* (between) neurons; both manifest as topological errors. Collisions between neurons are particularly harmful because they can create connected segmentations, erroneously including dozens or hundreds of other nearby cells. To make up for poor software, scientists assume they must lower the chemical labeling further. While this does lower the frequency of such fused clusters, this approach is short-sighted since it lowers the neuron yield per animal. Lower reconstructed counts per brain mean vastly longer and more expensive experiments. Scalability is of principal importance to handle the power-law distribution of cells and their inevitable clusters.

Poor accuracy can also arise from problems at the image level such as artifacts, thresholding, normalization, pixel-level noise and anisotropic blurring. Neural network approaches tend to have a low-frequency bias and regularization techniques tend to increase dimensional blur instead of correcting for it as demonstrated in NeAT [41]. Neural functions also tend to lack continuity on thin signals as mitigated by Lipschitz regularization [28]. Additionally, CNNs can be prone to oversegmentation and path collisions since there is not an adequate loss for the topological effects of detection. For example, though less fine than neurons, the thin struts of chairs in ShapeNet [15] often create collisions and breaks.

Tracing from seed locations to their connected branches can be conducted by propagating a wavefront spatially across a grid, in a general pattern known as the Fast Marching Method (FMM)[42]. Though FMM is a core part of the top neuronal reconstruction methods discussed in the skeletonization section, it is inefficient both in time and space. Brain volumes are large (many Terabytes), highly sparse (less than 1% foreground) and individual neurons can span the whole volume. When FMM computes these images it leads to extremely low arithmetic intensity and runtimes dominated by streaming dense image regions from disk repeatedly[32].

Very similar in process to Dijkstra’s algorithm, FMM leverages a priority queue to choose its next visited location, which dictates its *O*(*nlogn*) time complexity. The FMM computation differs from Dijkstra’s in that it stores each path distance across the whole domain grid, which makes it *O*(*n*) in space. The Fast Sweeping Method (FSM)[52] improves the time complexity to *O*(*n*) by *sweeping* updates in diagonal planes. Refer to [52] for an explanation of how the diagonal access pattern converges to the same solution as pulling updates from a queue. The *parallel* FSM algorithm[53] leverages the fact that dependencies are independent along these planes. FSM is an excellent basis for implementation of applications since it is both parallel and has the favorable algorithmic benefits of sweeping.

The OpenVDB library[35] provides methods and data structures for spatially sparse volumetric data (Figure 3). However, modeling the domain in a dense fashion is inefficient. Much of the advantages of voxel grids–an almost infinite domain size and lower computation and memory–stems from their innate concept of a sparse active foreground set. VDBs, like other similar sparse data structures such as B+ trees[8], trade-off memory footprint and access speed, due to hierarchically compacted data at various resolutions. Due to an optimized hash map, they retain average case *O*(1) even for random access, updates and deletions. Higher tree depths yield better compaction of sparse signals at the cost of indirection steps (cache misses) at each level. Following from the observation that accesses are generally spatially coherent, VDB reads and writes are mediated by an *accessor* structure which leverages coherence when finding new elements via a bottom-up search from a cached set of recent visits.

**Fig. 3.**
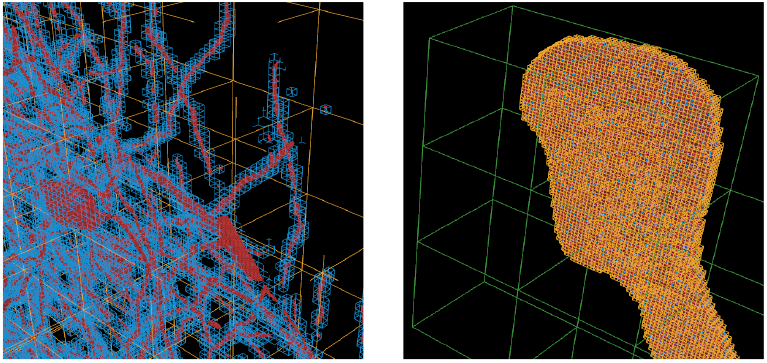
A visualization of the various spatial levels of granularity of VDB grids on a sparse neuronal slice of data. The outline of the cubes are highlighted to get a sense of the 4 scales. The sparse working set (*W*_*s*_) of foreground voxels is shown in red. Active leaf node regions–those that contain a foreground voxel–are outlined in blue. Active internal nodes regions are outlined in yellow and active secondary internal nodes in green. Empty leaf nodes and internal nodes have no outlines and have tiny memory footprint overheads. Refer to [35] for a further explanation of this terminology and VDBs.

The drawback of OpenVDB is that operating even on the whole sparse working set (*W*_*s*_) is excessive. For medical imaging volumes such as brains, these methods would consume an estimated 40 TB of peak RAM which is currently beyond what is feasible on a single workstation. We explain these working set (*W*) definitions and how to circumvent these issues in Section 3.

### D. Skeletonization

Abstracting segmentations into ball (with radii) and stick (connections) models (see Figure 4) is known as *skeletonization*. This is useful for (1) efficient proofreading and interaction with labels and (2) analysis of morphological and topological features and errors. Neuroscientists primarily model cells and their topological branching patterns with these skeletal graphs. More specifically, they use *trees*: each node in the graph has a single unidirectional edge which traces a path back to the tree’s seed (the cell body) [14]. This process is bottlenecked by scalability because of entrenched domain-specific methods.

**Fig. 4.**
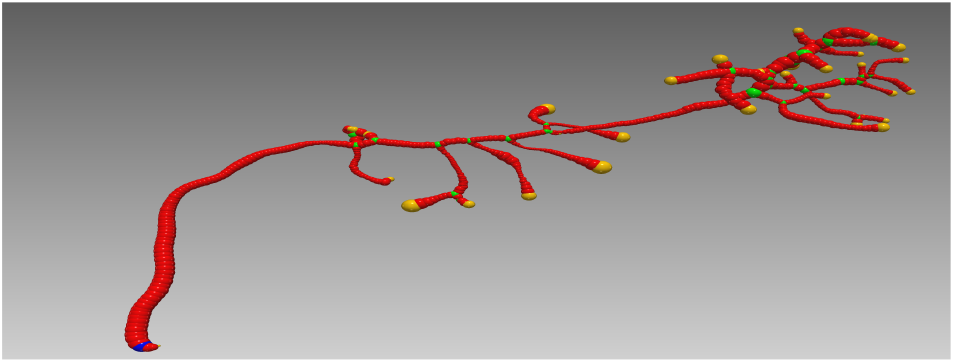
The ball (spheres) and stick (obscured connections) abstraction that neuroscientists work with. Note that this is a spatially embedded tree graph rooted at the seed which is the blue sphere at the bottom left corner. Red spheres are path nodes, yellow spheres are termini/leaf nodes and green spheres are branch nodes. Refer to [14] for a further explanation of tree graph (SWC) terminology.

There are hundreds of semi-automated (requiring human correction) implementations specialized for neuronal reconstruction [1, 30]. However, these methods lack *scalability*– the ability to handle large image regions. This limits their usage to (1) neuron types that are highly localized, (2) small hand-cropped volume sizes, and (3) neurons with little or no collisions. To our knowledge, there is no other method for reconstruction that does not either manually or automatically crop the neurons arbitrarily for performance reasons. If these alternative methods did not do so, neurons with collisions would be lost due to timeout or excessive reconstruction time. The majority of other algorithms also operate in 2D due to runtime concerns and maintain both a heavy object-oriented view of the data while also consulting the dense representations of the image volume. This coupling results in extremely high memory footprints and low arithmetic intensity as we demonstrate in Section V. Enabling full concurrency and exploiting the cache hierarchy through causal and spatial sparsity nullifies these issues, but only if every working set can fit entirely in memory (see Section V-A for the specifics of our system).

Based on benchmarks comparing numerous neuron reconstruction methods [50][31], APP2[49] has the lowest runtime and resource usage and the best algorithmic complexity. APP2 also placed in the top 5 highest accuracies, with Neutube [20] placing first in [50]. Neutube [20] as well as many high-performing reconstruction methods [4] employ a fastmarching algorithm similar to APP2. Such methods are all of *O*(*nlogn*) complexity. As mentioned because of excessive runtimes, a common practice is to manually crop around a neuron which can create cutoffs, artifacts, and intensive human intervention. For example, a recent prepublication [29] with similar pipeline goals only allows a maximum crop size of 512×512×256 and applies APP2 on these crops due to performance constraints. Such small sizes are only suitable for a subset of neuron types. APP2, like other available iterative reconstruction methods, is only single-threaded. This is why large-scale applications must start multiple APP2 programs for each seed to enable some concurrency [25]. We compare to APP2 for performance and accuracy metrics.

### E. Application Pipeline

Neuroscience data sizes tend to increase at the rate of 1000× every 4 years. This explosion in the acquisition of unstructured data creates problems which we explore below.

As shown in Table 1, Mouselight [48] is an interactive visualizer of large volumes. Its respective memory footprint reflects the dense volumes size of the images we tested. It relies on a human-guided tracing of neurons which takes several hours per neuron. APP2 [49] reflects the max sizes we could complete on a 4 TB system with our application (in single threaded mode) to auto-crop the region to its bounding box. The commercial software Imaris has an alternate cropping feature but the process is manual per neuron and also fails on images this size. Our method, APP2 and an APP2 based pipeline [29] all trace neuron branches automatically, though they need subsequent proofreading.

**TABLE I.** We compare the maximum scale allowed for a single neuron and the memory cost associated (RAM) to 3 major related works. Our method is 67× larger in scale with 870× less memory than APP2 which is the most efficient alternative automated approach.

| Work | Max. neuron | Auto | Mem. GB |
| --- | --- | --- | --- |
| SEU pipeline [29] | $512\times 512\times 256$ | ✓ | 11 |
| APP2 [49] | $3788\times 5380\times 1995$ | ✓ | 2609 |
| Mouselight [48] | $13000\times 13000\times 16000$ | | 2704 |
| **Ours** | $13000\times 13000\times 16000$ | ✓ | 3 |

Applications in this space offer features in the categories of *creation, mastery* and *distribution*[23].

#### Creation

This involves tools for generating, editing or authoring assets. Much like the autonomous driving space, in microscopy, large-scale noisy 3D volumes are plentiful yet instance-level models (neurons), especially those that are corrected and labeled by type are limited. The diversity and uniqueness of scientific queries similar to artistic endeavors makes creating specific data even more valuable. We contribute an application that creates a variety of 3D representations. From raw images, we improve image quality, segment cells and output volumes, 3D models (meshes), skeletons and training labels (image window crops). We also provide the test suite *TreeBench* to establish the accuracy of these 3D assets. We rely on the existing tools Neutube[20], Fast Neurite Tracer (FNT)[21], and TeraFly[13]. All of these are volume-rendering GUI tools for either (1) proofreading our automated outputs or (2) creating neuron skeletons or seed labels from scratch (in the case of inadequate skeletal accuracy).

#### Mastery

This relates to utilities that facilitate discovery, search, reuse and understanding of data. Solving issues in scale brings new problems in the management of numerous outputs. Due to operating on whole brains, we generate hundreds of automated reconstructions per brain. As a result of their multiplicity, these reconstructions have diminishing marginal value unless they are automatically sorted according to their quality which has become untenable to do manually. We filter and tag outputs automatically based on quality.

In the creation of neurons, we use low-bias unsupervised techniques. Our aims are minimal: we seek that paths are continuous, do not collide and do not have unfaithful mutations. Generic transformations are more adaptable because the scientific study of morphological differences is subtle and can be corrupted by the influence of out-of-distribution training data. This allows the outputs to be generic and therefore more likely to be candidates for reuse in future scientific queries. Finally, a semantic understanding of cell bodies and their branches allows automatic tagging and labeling of images. This automation can help bootstrap future methods and more specialized experiments on unforeseen data types.

#### Distribution

Contributions in this category aid the dissemination and sharing of assets between groups. Mouse brain volumes are on the scale of several terabytes, thus sharing and storage are problematic. Since our end-to-end application is essentially domain-specific compression, whole brains have extremely low memory footprint representations (less than 2 GBs) which can be output at various stages, resumed and shared across computers or teams easily. We apply strict final passes of the skeletons to adhere to the community standards established in StdSwc[5]. This allows interoperability with outside or legacy tools and is a requirement before it can be added to the community database NeuroMorpho.org[2].

## III. Method

Computer vision tasks involving 3D geometry or volumetric data should leverage not only *spatial* but also *causal* sparsity. *Spatial sparsity* reduces computational demands by exploiting local structures in data. Segmented regions and their values, for example, tend to be spatially coherent within various scales. Languages [24], libraries [35] or applications [34] that leverage patterns in spatial distribution are an increasing necessity on simulations at scale.

*Causal sparsity* further decreases resource usage by dynamically growing a smaller working set as needed from key salient structures or regions. We term the set of these key features or elements *seeds* and refer to them as {S}. This approach separates a large-scene into *object-level* tasks where instance-level computation can exploit more temporal locality. Spatial and causal sparsity complement each other. Not only can they share data representations, but their locally restricted access and search patterns can offer best-in-class performance.

The core of our method can be broken down into 3 specializations of search:

1. *W*_*s*_ *scan:* traverse a dense working set *W* such as an image volume and choose a semantic subset W_s_.
2. {*S*} *scan:* find key elements termed seeds {S} within W_s_.
3. *W*_*c*_ *select:* iteratively apply a recurrence relation from {S} to select a causally sparse working set W_c_ as needed.

These patterns are particularly effective on data-intensive parallel applications. For example, this combination of spatial and causal sparsity dramatically increases the image scale that can be processed on a single workstation. In the context of this manuscript, the dense 3D cube (the whole image) is the original working set *W*. Refer to the grid diagrams (labeled 8-11) in Figure 5a. In these grids, the labeled regions are shown with white colored pixels–this is the spatial sparse working set W_s_. The blue colored regions are the cell segmentations that specify a causally sparse working set W_c_. Refer to Table IV for the element count reductions. W, W_s_ and W_c_ are three views of the same data. Their successively smaller element counts lead to empirical performance gains. Problems, especially those arising from natural data, often start with a dense working set W that can be compacted to a spatially sparse working set W_s_. W_s_ represents a set of relevant elements narrowed *spatially*, whereas W_c_ is the often much smaller dynamic set of values derived from *causal* selection. W_s_ scan can also minimize the representation or place it in a specialized data structure to suit later computation. Selective search allows algorithms to terminate early, thereby preventing unnecessary accesses and computation (i.e., *O*(*W*_*c*_) as opposed to *O*(*W*_*s*_)). For example, if regions of an image are not reachable from a seed and thus not *causally* linked then they are ignored (see the connectivity of the diagram in Figure 2).

**Fig. 5.**
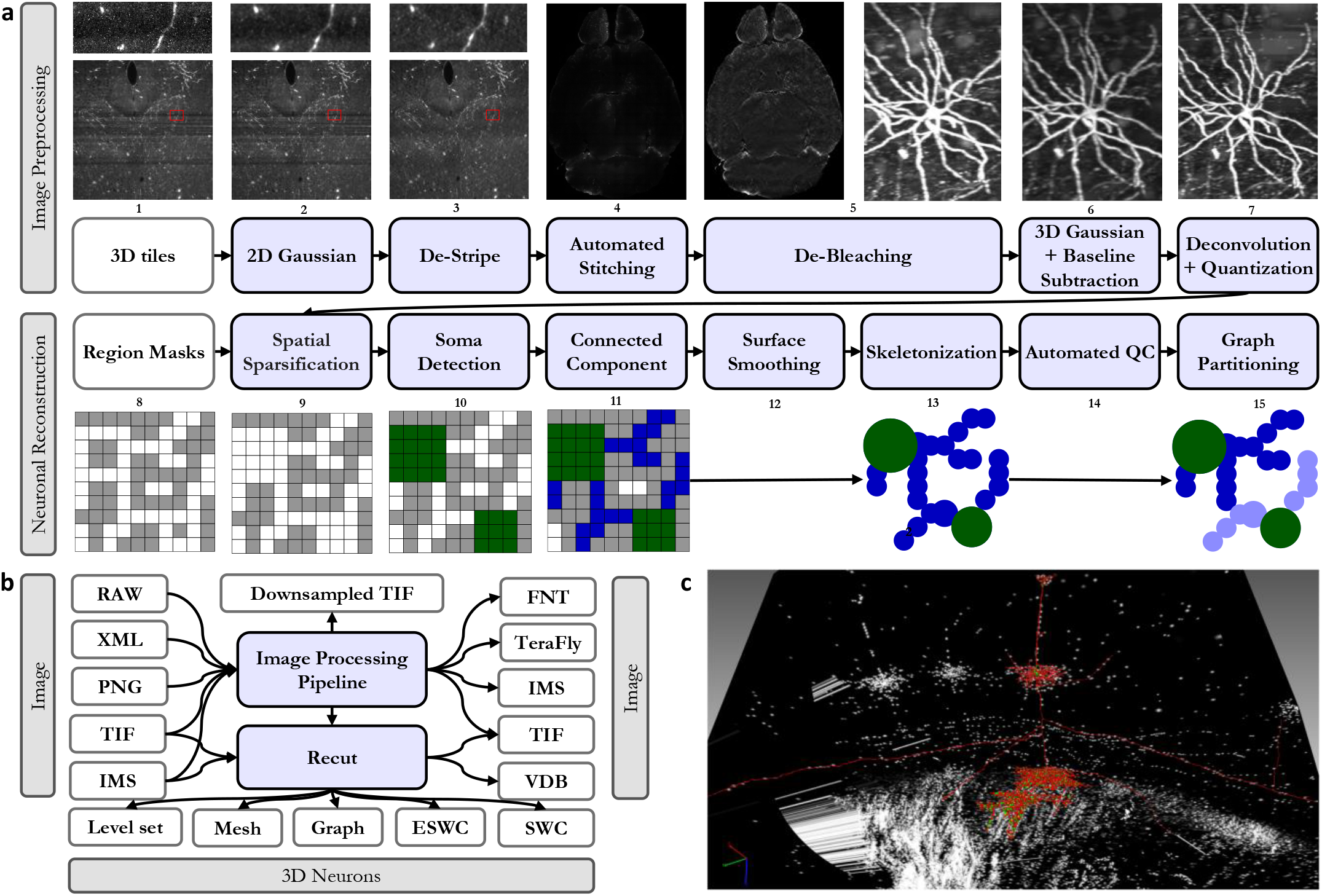
Detailed data processing pipeline towards the reconstruction: (a) We suggest a specific order in which the data needs to be processed to improve the quality of reconstructions. (b) Gossamer pipeline produces a wide variety of outputs for different needs. (c) A sample reconstructed cell that is visualized in neuTube software and is ready for human validation.

In this work, our example scan and select functions are of linear complexity which is essential for problem sizes with power-law distribution in scale. Scans are trivially parallelizable and are generally decomposed spatially, whereas selection is iterative but parallelizable across seeds.

Narrowing the working set is effective across varied graphics methods because data is often relevant mainly in contextual relation to seeds. This relevance can often be decomposed into a chain of computations leading back to a seed. Therefore, exploiting causality involves identifying seeds and allocating or weighting resources and computation according to their seed relation.

## IV. Implementation

We abstract these design patterns and apply them individually, on 4 example computer vision workloads, and in combination with a challenging application. Each example workload described below can be broken into scans and selections. At a more abstract level, each workload itself is analogous to one of these patterns, especially when viewed in a feedforward pipeline. Encoding is analogous to a W scan, seed segmentation to a {S} scan, cell segmentation, a W_c_ selection and skeletonization, a W_s_ scan. These scan and select patterns offer a neuron reconstruction method that is fully parallel, hardware-aware and entirely in-memory.

### A. Encoding

Image processing enhances the image. This allows continuing with a far more minimal view of the image while still retaining the structures of interest.

1. *W scan:* Destriping applies a 2D Gaussian and custom filters to remove camera artifacts and fine non-uniformities in the image tiles (steps 2-3).
2. *W scan:* Debleaching removes coarse non-uniformities in the stitched image (step 5).
3. *W scan:* Deconvolving improves the axial resolution and improve the brightness of the foreground signal (step 7).
4. *W*_*s*_ *scan:* Conversion traverses the image volume, placing voxels above a value in a spatially sparse data structure (VDB). W_s_ is the segmented mask of the image which holds all foreground voxels (step 9).

#### Destriping

Using 2D wavelet decomposition, the image is decomposed into horizontal, vertical and diagonal coefficients. The horizontal and optionally vertical coefficients are decomposed with 1D Fourier transform on the appropriate axes. The Fourier transforms are multiplied to a Gaussian notch filer with a desired sigma. Finally, using inverse Fourier transform and 2D wavelet recomposition the image is reconstructed.

#### Debleaching

First, the stitched image is destriped bidirectionally to correct regions that are directly bleached by the laser beam. Finally, a low-pass filter is applied to the image in order to capture the gradual change in the brightness of the background from the surface of the brain to its center that happens because of poor penetration of laser light. The image is divided by the filter to reverse the bleaching.

#### Deconvolution

In lightsheet microscopy, the image of a sphere appears convolved by an ellipsoid kernel. By knowing the kernel shape, the original shape of the object can be restored via inverse convolution operation.

#### Conversion

Due to the substantial increase in signal to noise ratio from the previous steps the image can be segmented by keeping pixels with sufficiently high intensity (brightness). To make conversion more robust across different brains, we use a default desired foreground percent and compute a hard threshold value that most closely matches the foreground percent. We do so with quicksort, the only place in the pipeline outside of the skeleton standardization passes that is not of linear algorithmic complexity.

### 1) Parallel Implementation and Software Pipeline

Scan steps are embarrassingly parallel in nature. All stages at the encoding level must operate on the extreme size of the dense W volume (see Tables II and IV). Therefore, we parallelized all W scan steps across multiple GPUs and CPU cores and in so doing developed several programs with significant advancements in scalability.

**TABLE II.**
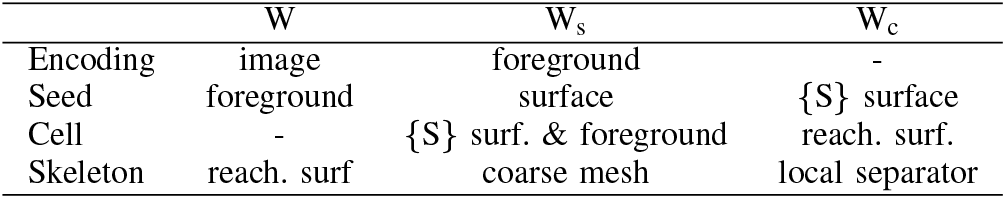
Since each stage progressively shrinks the working set, *W, W*_*s*_ and *W*_*c*_ take on different meanings at each stage. The derived outputs of one stage are often the inputs to a later stage.

1. We implemented a multi-GPU accelerated version of Pystripe [44] that has a novel bidirectional destriping and Gaussian denoising algorithm as well as a faster parallel processing model.
2. Our software offers a multi-GPU accelerated version of TeraStitcher [11] image stitching software.
3. We developed a multi-GPU and parallel version of LsDeconvolve [9] – an open-source program for the deconvolution of lightsheet microscope images.
4. `process_images.py` is a script that automates the stitching process.
5. `conver.py`, is a high-scale parallel file conversion tool for the various formats shown in Figure 5b.
6. The FNT cube processor program can apply the destriping algorithm on the z-axis of FNT files.

Our conversion tool can read Imaris files, 2D TIF series, or XML output of Terastitcher to convert them to processed 2D TIF series, Imaris, TeraFly, FNT or MP4 video while applying the above-mentioned filters (destriping, bleach correction, baseline and background subtraction, 16-bit to 8-bit conversion), or resizing or downsampling images to a desired isotropic voxel size. Our stitching implementation can also optionally apply a destriping algorithm, a low-pass filter to the stitched image to remove coarse bleaching artifacts and can downsample the data. Finally, all the above-mentioned image processing tools are in-memory to reduce IO overhead and also have resume support and advanced error-handling mechanisms to reduce day-to-day operational challenges. See supplementary notes 1 to 8 for more information.

### B. Seed segmentation

It is critical to first detect cell bodies (*seeds*, shown in green below) from brain images (black and white grid) to further segment each neuron’s connected branches. We do so with the following (steps 8-10 in Figure 5a):

1. {*S*} *scan:* convert the segmented mask VDB from the previous thresholding step into a VDB which stores voxels only at the surface of the foreground *W*_*s*_. {S} is then the surface voxels of W_s_ (step 8).
2. *W*_*c*_ *select:* seeds are segmented by successive region growing steps outwards and inwards from {S} (step 9-10). W_c_ is the resulting segmented cell body surfaces.

Cell bodies are large and spherical and can therefore be segmented by spatially expanding and contracting the foreground regions. The images are chemically labeled only on the cell surface due to the usage of a novel genetic technique [46] specially designed to improve reconstruction on thin, finegrained and long-range structures. Since the signal inherently describes *shells* [46], it lends itself naturally to the sparse level set methods provided in [35] which our application exploits effectively. To fill hollow cell bodies we perform morphological dilation. Since dilation expands the sizes of signals we apply equal and opposite erosion. Together this is known as morphological *closing*. To remove the cell branches which are thin and tubular we employ erosion followed by an equal amount of dilation, which is referred to as *opening*. This recovers seed regions whose geometries are relatively unchanged. Refer to the grid diagrams of Figure 5. The output of seed segmentation is the green regions. We use a single-threaded implementation of connected component analysis provided by the OpenVDB library to separate all somas into a set of instances.

### C. Cell segmentation

We can segment entire cells (step 11 Figure 5a) by computing reachability (blue regions below) from the known cell bodies (green) detected in the previous step 10 (i.e., {S}) with:

#### Algorithm 1

Seed segmentation

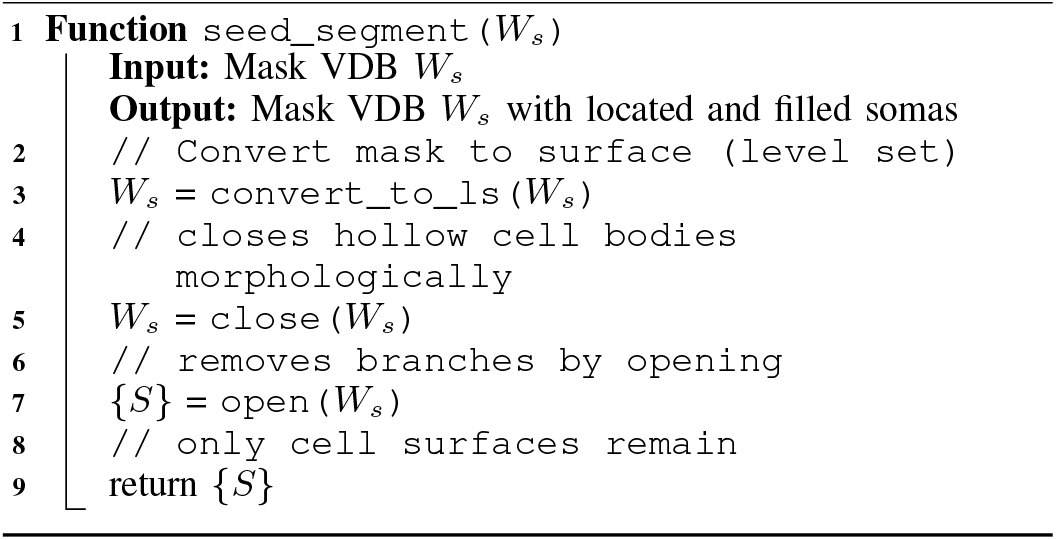

1. *W*_*c*_ *select:* from cell body surfaces, apply region growing outwards by consulting W_s_–the segmented image mask– to connect all reachable branches W_c_ (step 11).
2. *W*_*s*_ *scan:* traverse the grid to prune voxels whose values indicate unreachability. The kept voxels with a valid reachable distance compose a new W_s_.
3. *W*_*c*_ *select:* from the new W_s_, separate disjoint cells in a connected component analysis. This yields all instance segmentation surfaces separated into individual neurons.

We employ the fastsweeping method in [36], but rather than use it globally we start it from the key semantic areas of interest: the cell bodies from the previous step, and allow it to naturally terminate once its computed distance on all *reachable* voxels. In the grid at right, notice how the reachable voxels have a distance computed whereas the unreachable voxels are empty indicating an invalid value that is discarded with a subsequent parallel scan. Fastsweeping is generally applied on dense 2D or 3D regions (i.e., *W*).

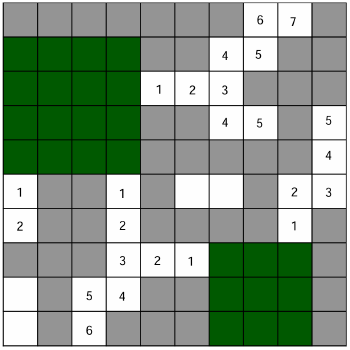

The VDB library improves on this with a sophisticated sparse implementation. This is a key advantage to VDB’s sparse methods, although one can have an almost infinite domain size, performance costs only reflect the sparse voxel count *W*_*s*_. However, when *W*_*s*_ is still too high, as in our case, application on all of *W*_*s*_ is a drawback. Instead, we start the selective search only from known cell bodies. This further improves performance and reduces peak memory usage since only the reachable voxels (e.g., *W*_*c*_) are accessed during the *W*_*c*_ select. Tables II and IV enumerate the differences between *W*_*s*_ and *W*_*c*_. Table IV in particular indicates the reduction in computation and memory of our approach. In addition to an implied contiguity (reachability), fastsweeping also computes a maximum distance from any cell body surface. This is employed to finally prune the voxels that have a default (maximum distance) value. Although this approach still entails a final prune over all of *W*_*s*_, pruning is a scan which, unlike fastsweeping’s selective search, has perfect parallelizabilty, minimal computation and memory requirements and almost no overhead.

During segmentation, we also optionally employ morphological closing of the branches. Whereas the foreground percent parameter merely keeps or rejects different values in the image, closing is a rather biased mutation of the morphology. In theory, it mitigates path breaks at the cost of path collisions, a relationship we explore in the results. A higher foreground percentage may be preferable for faithful surface reconstructions however closing is tested as an alternative method to keep the global *W*_*s*_ low.

#### Algorithm 2

Cell segmentation

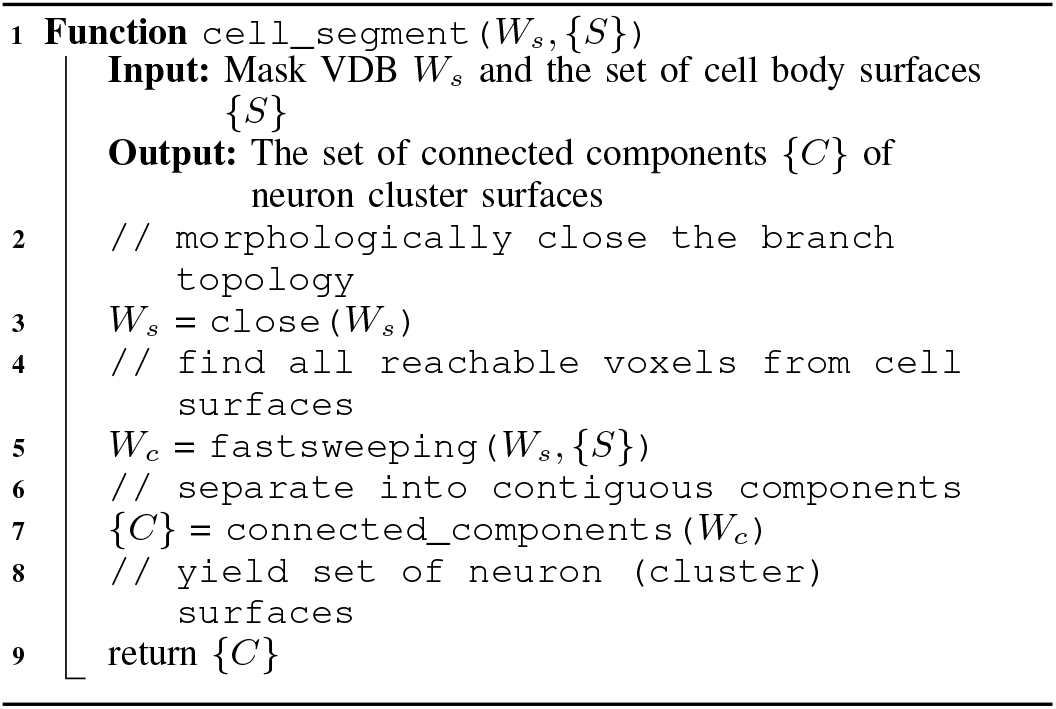

### D. Skeletonization

Skeletonization of the segmented cells creates a compact ball and stick representation–a spatially embedded graph with radii information (steps 12-15 in Figure 5a). Reference [18] gives a formal definition of *curve skeletons*. This allows neuroscientists to effectively study topological features. We do so via:

1. *W*_*s*_ *scan:* traverse the set of surface voxels and create W_s_–the spatially sparse mesh that retains topology (step 12).
2. {*S*} *scan:* using heuristics, choose a set of seeds from W_s_ (step 13).
3. *W*_*c*_ *select:* from each seed, iteratively search for a separator by expanding a sphere and adding encompassed vertices (step 13).

We follow the local separators (LS) skeletonization technique [7][6] but add an initial W_s_ scan to prevent high runtime at high resolutions. Due to multi-level coarsening of the surface mesh and projection as explained in [6], skeletonization achieves *O*(*n*) complexity. LS is particularly scalable on tubular signals especially with mesh (surface) representation–a volumetric representation would have less favorable performance.

The LS has a rather subtle advantage in reconstruction fidelity over other methods: it is topology-native. It can recover essentially all, even tiny, highly fine-grained protuberances and bumps while still capturing topology at vastly larger scales. This becomes apparent when switching to a 3D surface representation, where we find that neurons have small spines and projections on the surface. These details are often not visible in existing volume rendering tools unless the volume is binarized and projected in 2D. This ability stems from LS employing the *select* pattern until the wavefront (neighbor vertices of those visited so far) splits into more than 1 connected components. Our approach decouples anisotropy, resolution and scale differences in structures irrelevant from the skeletonization process. Unmanaged, this ability would come at a cost to runtime.

Since neuroscientists historically are only interested in the broader topology of branches and not surface features we remove these before skeletonization via, an optional, surface smoothing of the mesh. Since this mesh pre-coarsening step has the added benefit of a net gain in speed (about 44×), it follows that we use the simple and fast *light-edge matching* approach which is also employed in the LS work. This method guarantees not to introduce new path breaks or path collisions (constant number of connected components), but can produce artifacts related to smoothing. It operates by a user-specified number (coarsen steps) of successive linear scans.

Mesh coarsening allows us to reconstruct only the desired topology of neurons while remaining accurate and performant regardless of the resolution of the image. Even higher resolutions would benefit further from this option, therefore coarsening steps is a runtime parameter determined by default by the image’s voxel size. APP2 does not use a mesh intermediate representation which makes its performance and resource usage highly sensitive to resolution.

#### Algorithm 3

Skeletonization

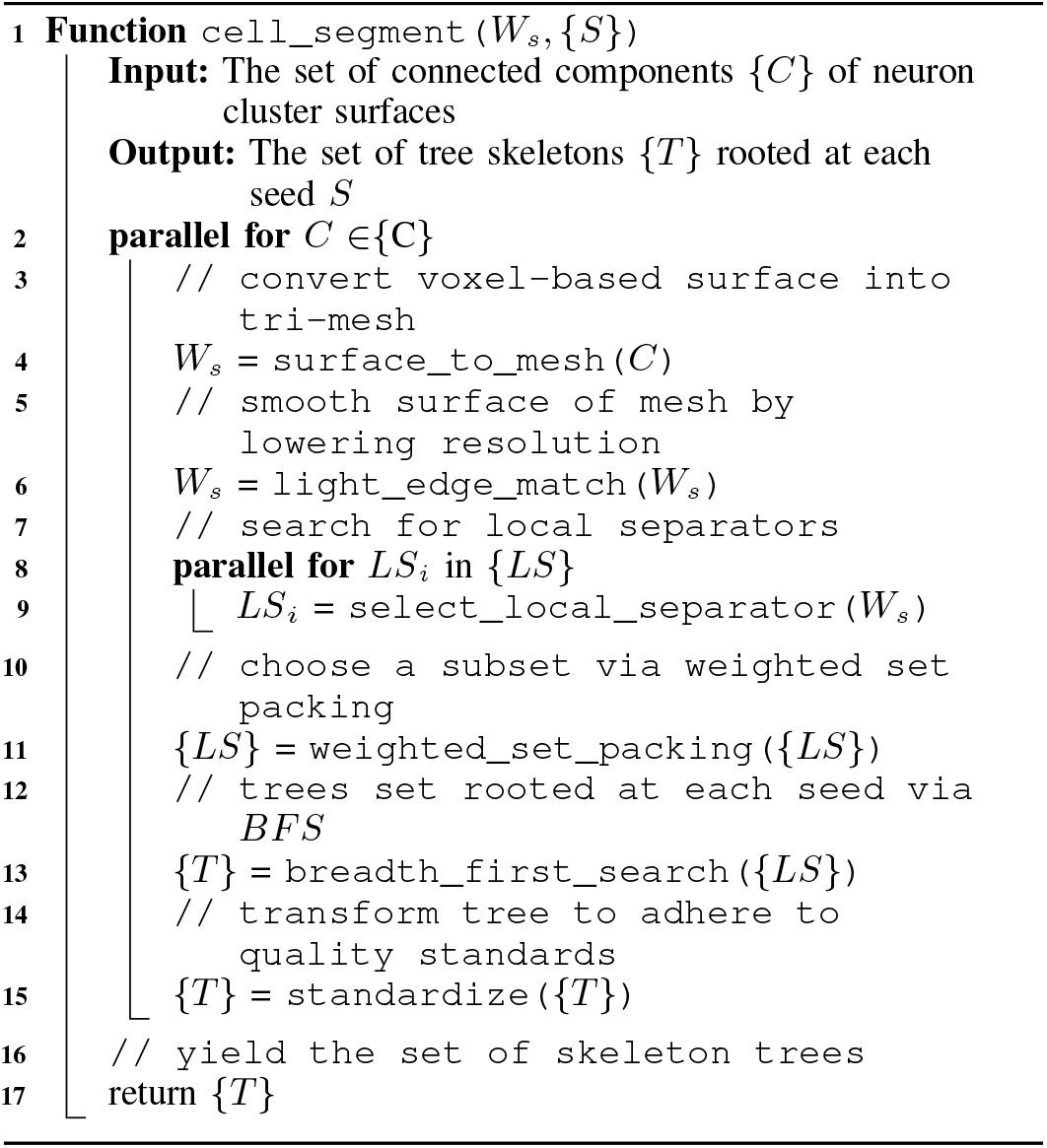

## V. Experiments and Results

### A. Data, Setup and Code Availability

We intentionally implement for a single workstation, as that could allow for the application to be deployed on the same microscope computer that acquires raw images. For massively compressive pipelines such as this, we advocate for computation on the *edge* as opposed to *distributed* due to the prohibitive network cost of the latter. Therefore for these workloads, we recommend a single workstation with 2× AMD EPYC 7763 (128 cores/ 256 threads) CPUs, 4 TB RAM and 8× A100 GPUs. The large memory capacity allows data sizes of 1 whole mouse brain at 1um^3^ resolution to be processed entirely in-memory for all tasks. We also test an equivalently sized volume at the higher .4um^3^ resolution to test accuracy effects. The 1um^3^ brain has 374 gold-standard neurons of a variety of neuron types with which to compare. Note that only a highly sparse subset of neurons (500-5000) per brain are labeled and only on the cell surface due to leveraging a novel genetic technique[46]. Some with far-reaching branches that span a large portion of the volume. In contrast, the 181 labeled neurons used for the .4um^3^ dataset are local and bush-like which allows us to compare to APP2. Both image volume dimensions are approximately 13000×13000×16000 voxels, thus the higher resolution image only captures approximately half of a brain.

We manually proofread all our automated outputs twice using Neutube [20] or Fast Neurite Tracer (FNT) [21] to establish our ground truth skeletons for comparison. We embed an APP2 implementation into our codebase to lend it automated seed segmentation and windowing around the neuron for a fair comparison. We also supply APP2 with our foreground percent algorithm since its native segmentation method, a raw background threshold value, fails completely.

We tested various foreground percentages with TreeBench and found metrics tend to peak around .8%. Unfortunately APP2 can only complete a single neuron (the smallest one) at this density, therefore all APP2 runs are at .4%. These parameter choices illustrate how entangled performance and accuracy is for poor scalability algorithms. Code for the reconstruction and image preprocessing pipeline is available at https://github.com/UCLA-VAST/recut and https://github.com/ucla-brain/image-preprocessing-pipeline respectively.

### B. Encoding

#### 1) Empirical scalability and performance

For our images, destriping and debleaching take about a half a day to complete. Deconvolution’s runtime is also about a half day. As illustrated in Figure 7, conversion of the dense image W to the spatially sparse VDB representation W_s_ takes 15.5 hours on a single core and 21 minutes on 128 cores, which is about a 44× increase in speed. Before the VDB conversion the working set was over 3 orders of magnitude larger than the spatially sparse working set W_s_ used for the remainder of seed segmentation (Table IV).

On a Windows workstation equipped with 2x Quadro RTX A6000 GPUs and 2x Xeon Platinium 8260 CPUs, GPU acceleration improved the destriping speed from 10 image/s to 73 images/s in our testing.

#### 2) Accuracy

We test the efficacy of preprocessing steps in 3 ways. We confirm qualitatively that known artifacts are removed after each stage (top row in Figure 5a). Compressibility is a proxy for signal-to-noise ratio. Figure 6 shows that each stage other than deconvolution improves the compaction ratio of the image. Deconvolution sharpens the image which can exacerbate high frequency noise or surface details hence the slight increase in size. Finally, we test each encoding stage with respect to reconstruction accuracy as discussed in Table VIII and the ablations section.

**Fig. 6.**
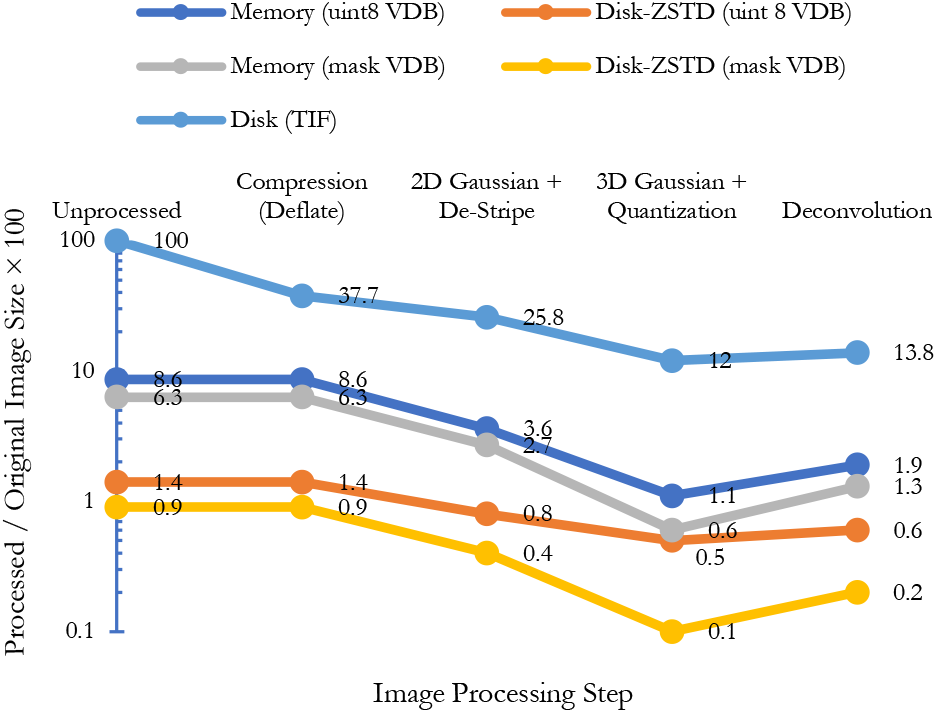
The effect of each image processing on image size both in memory and disk. Image processing steps and unstructured data (for example, TIF format) do not have any impact on memory usage but they can improve the data compressibility on disk. VDB format on the other hand can dramatically reduce both memory and disk usage. (a) Unprocessed TIF format occupies 100 percent of the disk and memory. The same data after thresholding and sparsification occupies less than 10 percent of memory and about 1 percent of the disk using a disk-assisted compression method. Interestingly, different stages of image processing increase the data compressibility not only for VDB files but also for compressed TIF files by removing noise and fluctuation in the signal. Using mask-VDB, the memory footprint will be a mere 1.3 percent for reconstruction in our pipeline.

**Fig. 7.**
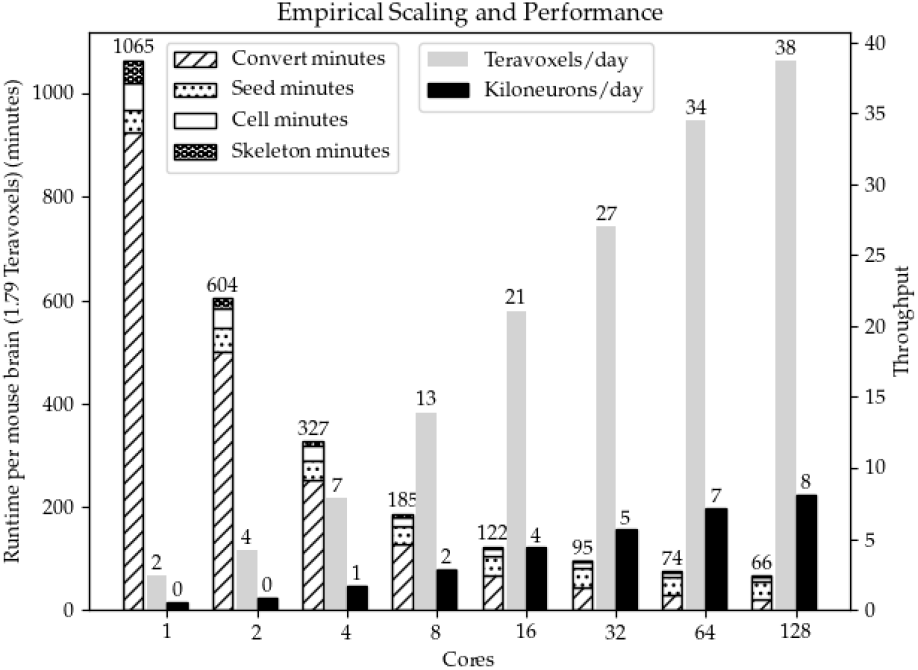
Aggregate runtime (left) and throughput (right) of the example tasks. Though scaling is sub-linear, added cores still yield significant speedup. More importantly, additional cores allow each task a higher peak memory ceiling which is critical at these scales.

### C. Seed segmentation

#### 1) Empirical scalability and performance

When fully parallelized, seed segmentation lasts roughly 35 minutes per brain. This is longer than the other workloads VDB conversion, cell segmentation and skeletonization combined (32 minutes) due to the single-threaded connected component analysis at the end of seed segmentation. The other two portions, morphological closing and opening, only take 48 and 5 seconds respectively.

#### 2) Accuracy

Automated seed detection has high recall as seen in Table III; it can recover .94 of true positive (TP) seeds. However it produces a high proportion of false positives (FP); only .06 of predicted seeds were true positives. We explain subsequent work to mitigate this behavior in the discussion.

**TABLE III.** Seed segmentation coordinate accuracy with 1 radius width tolerance for 1 whole mouse brain imaged at 6x 1um^3^ resolution. Recall is the fraction of true positives (TP). Precision is the fraction of false positives (FP) to TP. The sum of TP and FP represents the total automated seed and cell output count.

| Type | Recall | Precision | F1 | Count |
| --- | --- | --- | --- | --- |
| Auto | .94 | .06 | .12 | 8953 |

### D. Cell segmentation

#### 1) Empirical scalability and performance

Fastsweeping completes in under 9 minutes and cellular connected component segregation requires 23 seconds. Computing reachability from known seeds narrows the working set by almost 6× (Table IV).

**TABLE IV.** The Working set element counts for the dense image volume W, the spatially sparse foreground W_S_ and the reachable cell segmentations w_c_. The FACTOR COLUMN SHOWS THE SIZE INCREASE RELATIVE TO W_C_. W_S_ is roughly an order of magnitude larger and W is 4 orders of magnitude larger.

|  | Gigavoxels | Factor | Meaning |
| --- | --- | --- | --- |
| $W$ | 1790 | 10,400x | Dense image |
| $W_s$ | 1.01 | 5.87x | Foreground VDB |
| $W_c$ | .172 | 1x | Reachable from seeds |

#### 2) Accuracy

Fastsweeping is a simple heuristic that we use indirectly to find regions contiguous to a seed. Its utility depends on the continuity of the image after thresholding and conversion to the VDB representation (W *→* W_s_). In other words, it still suffers from path breaks or collisions if they remain in the converted image. Below, we examine the downstream effects on topology to best assess its efficacy.

### E. Skeletonization

#### 1) Empirical scalability and performance

Observe the empirical results in Figure 8: our method uses 3 orders of magnitude less memory per neuron than APP2. The explanation for this difference can be traced back to table IV. Per neuron, APP2’s is forced to operate on *W*, which is at worst 4 orders of magnitude larger if a neuron spans the whole brain. In contrast, our method is on the order of *W*_*c*_ both in time and space at the time of skeletonization. In addition to reading and retaining the dense 3D bounding box (crop window) of a neuron, APP2 also densely allocates several scratch pads, also on the order of *W*. On our system with 4 TB of memory, the largest neuron that APP2 could reconstruct (right most red cross) was of size 3788×5380×1995. This largest cluster took about 2609 GB of peak RAM. Our method can operate on a neuron or many neurons simultaneously that span the whole brain (67× larger than APP2) while also using 870× less memory. Skeletonization of 374 seeds takes under 2 minutes fully parallelized and under 43.5 minutes for 1 core (see Figure 7).

**Fig. 8.**
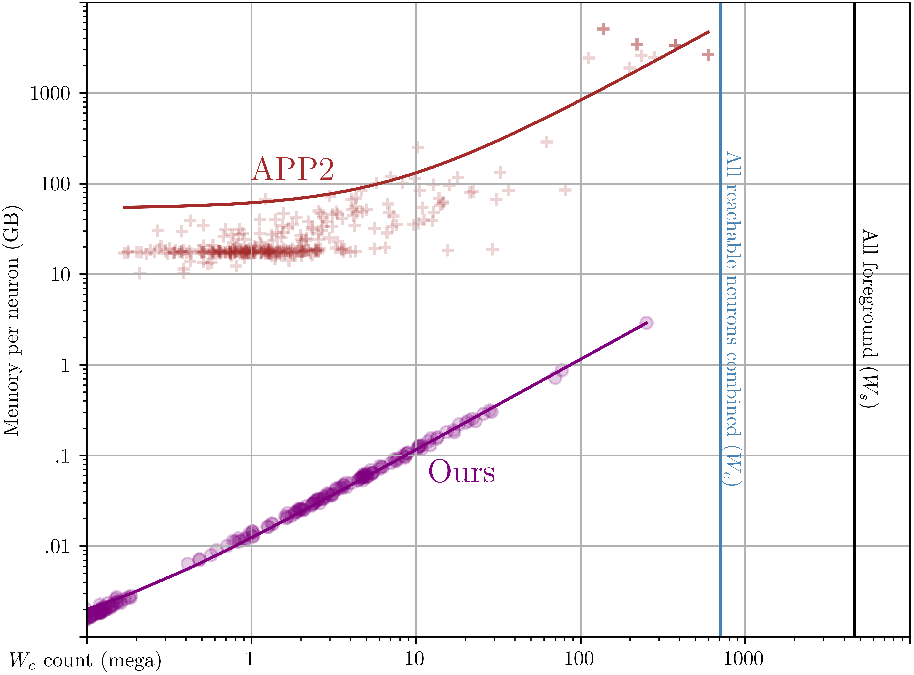
Crosses (APP2) and circles (ours) compare the memory allocated during of skeletonization of individual connected components (ideally 1 neuron). APP2 takes roughly 1000× more memory per neuron which is problematic with large clusters with intersections.

Although APP2 is the fastest and most scalable alternative method for single cell reconstruction, our method can skeletonize even the largest clusters in seconds where APP2 takes over a 100 minutes. Surface coarsening is largely responsible for this speedup since it allows reconstruction at a user-specified resolutions. We use 2 or 4 steps of coarsening since its more suitable for recovering topology (as opposed to surface structure). Note that APP2 pruning must approximate only in 2D due to performance, whereas our methods are all natively 3D. We summarize the differences in Table V.

**TABLE V.**
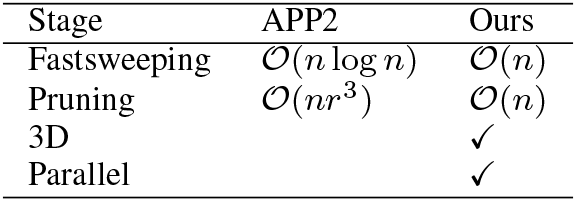
A comparison of the two major stages of skeletonization where *n* indicates the total working set count and *r* indicates the radius distance of a particular node.

### 2) Accuracy

DIADEM [22] is the gold standard comparison tool for neuron reconstructions largely because it has substantial correlation with both 1) proofread times and 2) expert subjective opinion on automated quality. DIADEM is a strict tree topological similarity score which yields 1 for a perfect match. For each ground truth skeletal node DIADEM checks 3 criteria:

1. Is there a *matching* node in the test skeleton within a Euclidean distance?
2. Do the potential node matches from step 1 have a corresponding *matching* ancestor node?
3. Is the path distance between node and ancestor and potential match node and its ancestor roughly similar?

Step 1 reports the basic recall or completeness of the skeleton. Step 2 prevents nodes on different or incorrectly directed branches from counting as a match. Step 3 further lowers the chance that nodes with a correct ancestor but a different path back to a seed get counted as a match. The distance threshold is dataset specific and is determined by the diameter of the largest branch point of all gold standard skeletons. For both datasets tested this value is set to 8um.

DIADEM is a good starting point for understanding the tree accuracy, however its largest drawback is a lack of transparency. To compensate, we isolate basic properties that correspond to known error types deduced during proofreading. Each of these topological qualities (see Table VI) helps explain DIADEM’s aggregated score. We term this suite, along with DIADEM, TreeBench. Its implementation is available along with the rest of the example benchmarks.

In tables VI, VII and VIII, topology indicates the DIADEM score which is an aggregate tree similarity metric. Recall is the fraction of ground truth skeletal nodes that have a matching node in the test. Branch and leaf also report the proportion of branches and leaf node matches respectively. Direction indicates the proportion of only the *matching* branch nodes that are also oriented in the correct direction which is why the score is often higher than the branch score. Precision is the fraction of test nodes that correspond to a node in the ground truth. Yield gives the proportion of automated neurons that had a topological accuracy within 2 standard deviations of the mean. Count illustrates the number of input seeds (cell bodies) used. The topology score reflects only the successful reconstructions, whereas all other scores aggregate errors across both successful and failed reconstructions.

When we adjust our mesh resolutions such that our runtimes roughly match those of APP2 per neuron, our yield is 78% higher (Table VI). We lend APP2 our foreground percenting algorithm (at the same .8%) for this comparison since it most closely matches the thresholding used by [29] another recent pipeline. APP2 accuracy and yield are far lower due to lack of the image processing steps of destriping, debleaching and deconvolution.

**TABLE VI.** Skeletonization accuracy for 1 whole mouse brain and 1 half brain imaged at 1um^3^ and .4um^3^ voxel resolution respectively. Our method at the higher resolution has the best yield indicating that even larger scales will have better accuracy with less proofread effort. The aggregate topological score is equal between our method at various resolutions but much higher than app2. The higher resolution is more effective at capturing fine-grained paths leading to more complete (recall) overall skeletons and especially termini (leaves). Our method at the lower resolution is able to resolve branch positions and their direction and the overall skeleton more precisely though that may be because it has a smaller, but more correct automated yield set of neurons.

| Voxel Res. | Ours |  | APP2 |
| --- | --- | --- | --- |
| | $1\mu\text{m}^3$ | $.4\mu\text{m}^3$ | $.4\mu\text{m}^3$ |
| Topology | **.74** | **.74** | .30 |
| Recall | .52 | .56 | .06 |
| Branch | .60 | .58 | .06 |
| Direction | .97 | .92 | .24 |
| Leaf | .28 | .39 | .03 |
| Precision | **.88** | .76 | .44 |
| Yield | .86 | .92 | .14 |
| Count | 374 | 181 | 181 |

**TABLE VII.** Treebench scores with several key parameters changed. We compare each ablation to the second column of Table VI (ours at .4um^3^) which has the default settings of 2 steps of surface coarsening, foreground percent of .8 and 4 steps of morphological closing. Each of these columns changes one of these parameters.

|  | Coarsen<br>4 | Foreground (%)<br>.4 | Close<br>1 |
| --- | --- | --- | --- |
| Topology | .67 (-Δ .07) | .69 (-Δ .05) | .72 (-Δ .02) |
| Recall | .53 (-Δ .03) | .50 (-Δ .06) | **.60 (+Δ .04)** |
| Branch | .46 (-Δ .12) | .58 | **.63 (+Δ .05)** |
| Direction | .93 (+Δ .01) | **.98 (+Δ .06)** | .97 (+Δ .05) |
| Leaf | .32 (-Δ .07) | .32 (-Δ .07) | .40 (+Δ .01) |
| Precision | .78 (+Δ .02) | .85 (+Δ .09) | .83 (+Δ .07) |
| Yield | .92 | .93 (+Δ .01) | .65 (-Δ .27) |

**TABLE VIII.** TreeBench scores for various ablations in image preprocessing. Again, we compare each ablation to the second column of Table VI (ours at .4um^3^) which applies all preprocessing stages. Each column is cumulative using the image processing step to its left. Anisotropic includes deconvolution (omitted with a double vertical line due to already being listed as the default in Table VI) but adds a final step of downsampling the z-dimension by a factor of 3 to probe the extent to which added information contributes to reconstruction accuracy.

|  | Raw | Destripe | Debleach | Anisotropic |
| --- | --- | --- | --- | --- |
| Topo. | .71 (-Δ .03) | .69 (-Δ .05) | .71 (-Δ .03) | .72 (-Δ .02) |
| Recall | .57 (+Δ .01) | .53 (-Δ .03) | .56 | .57 (+Δ .01) |
| Branch | .55 (-Δ .03) | .45 (-Δ .13) | .52 (-Δ .06) | .58 |
| Dir. | .93 (+Δ .01) | .92 | .90 (-Δ .02) | .91 (-Δ .01) |
| Leaf | **.43 (+Δ .04)** | .37 (-Δ .02) | .41 (+Δ .02) | **.43 (+Δ .04)** |
| Prec. | .84 (+Δ .08) | .82 (+Δ .06) | .81 (+Δ .05) | .81 (+Δ .05) |
| Yield | .67 (-Δ .25) | .91 (-Δ .01) | .90 (-Δ .02) | **.94 (+Δ .02)** |

Higher resolution can achieve better overall yield and less path breaks at the cost of more collisions and artifacts. Path breaks are the biggest loss in accuracy since a substantial fraction of ground truth skeleton nodes do not have a match in our automated outputs. Branch nodes tend to be more proximal to a seed and as expected have higher accuracy whereas leaf nodes which are the terminal points of branches have low recall. Of course, the global recall is in between both branch and leaf node recall, since it is an aggregate for all of a neuron’s nodes.

More complete neurons is often a priority when reconstructing neuron types that contain longer projections (axons). Reconstructing these neuron types is a key advantage since other pipelines can not capture them automatically at all due to performance. However, a collision or path direction error is much more time consuming to correct since it can involve deletion of a substantial proportion of the tree. In contrast, current proofreading tools are optimized to interactively fix path breaks.

### F. Application Pipeline Ablations

Surface coarsening lowers the granularity (vertex count) of the mesh surface, which also decreases the TreeBench scores as illustrated in Table VII. Although light-edge matching can not introduce new path breaks or collisions, the resulting lower resolution meshes do have the morphological artifact of shortened branches. The sock-like termini at the distal-end of cell branches have high curvature and tend to erode proximally (towards the seed) at 4 steps of coarsening. Very thin branches are more prone to this shortening, which causes losses in leaf and branch completeness, as well as overall recall, though to a smaller extent.

A lower foreground percent reduces the overall completeness and termini (leaf) recall due to breaks. Closing boosts the automated yield by a similar amount to the combined image preprocessing stages, however it only has these benefits when combined with these enhancements. However, decreasing the morphological closing (1 step instead of 4) improves individual scores of TreeBench which follows our observation that it is the most biased transformation we apply. For example, closing can move skeletal nodes to such an extent as to lower the matches. Branch points are particularly affected by closing, presumably due to erroneously joining the morphology of branch points further distally.

The highest yield occurs when applying all developed image processing techniques and additionally downsampling the z-dimension by a factor corresponding to the inherent point spread function of lightsheet microscopy. This smear in *z* is rather dramatic (about 2.5×) and deconvolution does not effectively remove it. It also limits the smallest resolution that skeletonization can place nodes and higher sampling leads to more accurate and natively smooth graphs.

## VI. Discussion and Future Work

Seed segmentation is effective at localizing and segmenting *potential* seeds, but performs poorly at distinguishing them from the background. This is likely because cell bodies are generic simple spheres that are difficult to discriminate from background noise after morphological closing. Adding a lightweight second pass classifier such as a multi-layer perceptron to cull false positives seeds could enable competitive accuracy. As it stands, humans must filter the seeds manually. Though these false positives can usually be batch deleted quickly with existing tools because they are clustered in unintentionally high saturation areas of the image.

It takes a similar time to compute reachability on a whole brain scale as it does similar methods like APP2 to complete fastmarching on moderate sized neurons. Cell segmentation is particularly fast because 1) it applies sparse fastsweeping 2) it starts with a large broadly distributed set of seeds which enables high concurrency with little ramp up and 3) unreachable regions are not accessed until the final prune step. Since the sparse working set W_s_ of cell segmentation is about 2 GB, it can stay in memory after conversion, and there are no disk read penalties of long-range connections. Other methods couple their algorithms with dense images, thus incurring reads from disk.

The local separators method [7][6] we used has linear time complexity. This is a major improvement over APP2 and Neutube’s *O*(*nlogn*) complexity which has compounding performance penalties when combined with the power law distribution of components. Although the number of seeds is generally larger than the thread count, some components are orders of magnitude larger. This leads to poor load balance, with some neurons taking much longer to complete regardless of thread count.

Our results suggest that even higher resolution would improve accuracy but scales much larger than 13k^3^, consume more than 4TB peak memory and are thus not possible to process with our method and reference system at this time. Still, we show that computational techniques can improve yield and accuracy with available resolutions.

### A. Distributed Approaches

One of our applications greatest strengths–residing in a single system’s main memory–points to a potential weakness: *out-of-core* image volumes. When the working set footprint is too large to fit in a single system’s memory, it necessitates either a sequential or distributed approach which we explore below. Gossamer has predictable resource usage estimatable by the working set size. This working set has a static memory footprint which we delineate from the *dynamic* memory. Though this peak memory can be several times larger than its static counterpart depending on the algorithm, it can be lowered by limiting the maximum count of in-flight threads. For example conversion of dense volumes to a sparse representation can complete on systems with far less memory by virtue of having much fewer threads active at a time (dictated by the hardware or by optional command line parameter). One can also run larger volumes sequentially and merge results via command line invocations and parameters. We apply this design aim–i.e., portability to limited resource platforms– throughout our pipeline.

We have decomposed the pipeline into either select or scan search functions. Scan functions can be simpler to automatically parallelize as they involve at minimum straight spatial block (cube) partitioning or, more generally for our stages, overlapping partitioning with halo regions. The large-scale U-net implementation in [32] follows this overlap then merge approach. Block partitioning has favorable (linear) scaling properties across nodes with the added overhead associated with overlap since different regions are entirely independent. The select pattern is less favorable to distribute across systems due to the excessive communication involved. A previous work[32] implemented block-based parallelism where threads pass messages to each other on halo regions across iterations of the algorithm on 3 example tasks including Fastmarching. This algorithm’s concurrency is similar to PageRank[37] and graph algorithms like NXGraph[16]. However, Fastmarching is a particularly problematic workload as it induces a surprising amount of communication due to the unknown causal paths in 3D space. With blocks of size 1024^3^, each cube reactivates on average 4 times until convergence. These reactivation factor increases with added signal density: a scenario where scalabililty is of particular concern.

Adapting the parallel implementations above for multinode concurrency in future work would have benefits and drawbacks. A distributed approach would have the advantage of the ability to scale to arbitrary large volumes. It would also be more cost efficient, as vertically scaled instances and hardware are disproportionately more expensive. We can look to the graph processing literature which has similar algorithms and both single and multi node schemes. Many single node implementations such as GraphChi[26], MapGraph[33] and X-stream[10] are more energy efficient with comparable latency and throughput as their distributed counterpart. Though stages like Fastmarching could be adapted such that messages were passed between processes on distributed nodes, this would greatly exacerbate the latency overhead at each iteration.

A larger concern than communication overhead is *load balance*–neurons are not evenly distributed spatially, even when considering the sparse set of foreground blocks of the VDB hierarchy. The best load balance, as exemplified by latter stages of the pipeline, occurs by partitioning not spatially but instead the set of *neurons* among threads. Of course, this requires first extracting neurons, which is done most efficiently with selective search. Recall that Fastmarching could not benefit from neuron-level load balance since it itself extracts neurons for the rest of the pipeline.

On a final note, main memory size has scaled impressively in recent years. In 2021, the maximum consumer grade server configuration consisted of a 2 CPU (dual socket) system with 256 total cores, 4TB of total RAM and 8 GPUs. 3 years and 2 CPU generations later, a single system can have 512 cores and 12 TB of RAM total. For these reasons all stages have been implemented for a single node with maximized computational resources.

### B. Conclusion

It is rather surprising that generalizations of search, when ordered effectively, can alter the scale in which neuroscience is performed. Competing methods employ highly integrated domain-specific heuristics that are coupled with dense image representations making them infeasible to parallelize. Our modular approach is more efficient and can be specialized for a particular representation and on growing datasets which allows it to gain in light of future data driven and supervised techniques.

### C. Contributions

We would like to thank the following authors and acknowledgments for their contributions to this work. We thank Tony Nowatski for his suggestions on an early paper draft. Muye Zhu provided the foreground percent algorithm and inspired the pipeline with the original MCP3D pipeline. Muye Zhu, Masood Akram, Chris Choi, Hongwei Dong, Chris Sin Park, X. William Yang, Zhe Chen, and Jason Cong advised on application and pipeline design and provided helpful discussions and feedback. Masood Akram helped evaluate reconstructions on gold standard datasets. Yuze Chi implemented an accelerated version of graph partitioning and fastmarching on FPGA. Chris Sin Park, Chris Choi and X. William Yang provided the mouse imaging data for the .4um lightsheet volume. Keivan Moradi and HongWei Dong provided the mouse imaging data for the 1um lightsheet volume. Chris Choi and Ming Yan led proofreading efforts related to the project and provided proofread soma locations for the .4um image. Ming Yan analyzed soma detection accuracy, provided scripts for overcoming manual steps in the pipeline including neuron filtering methods and designed a soma proofreading protocol. Brian Zingg and Chris Sin Park performed the surgeries needed to generate genetically labeled brains. Mitchell Rudd and Chris Sin Park performed tissue clearing and lightsheet imaging. Adriana Gutierez, Hyun-Seung Mun, and Qing Xue human-validated all reconstructions used in the manuscript. Sumit Nanda and Keivan Moradi, programmatically post-processed SWC files to prepare them for analysis. Sumit Nanda carried out preliminary testing of reconstruction output comparison with manual reconstructions using DIADEM metric Keivan Moradi designed and implemented image processing stages (encoding), wrote the corresponding portions of the manuscript and also second-pass proofread all 555 neuron reconstructions used in the manuscript. Karl Marrett designed and implemented application portions related to VDB, reconstruction and TreeBench, and performed the analysis and wrote the manuscript. This research is funded under NIH Grant No.: (U01MH117079-01).

## VI. Supplementary Material

**Supplementary Note 1** *PyStripe:* To better eliminate CMOS camera artifacts, we modified the PyStripe program to apply a small 5×5 Gaussian kernel with a sigma of 1 to raw images before applying the destripe algorithm. Empirically, the db9 wavelet and a sigma of 250 for 15x images and 100 for 4x images for foreground and background produced better results. Since the original PyStripe algorithm was adding artifacts to the edges of the image, we temporarily padded the images with the wrap method before destriping. The pad size is automatically calculated based on the destriping sigma. Note the pad size is equal to the half-height width of the Gaussian notch filter.

We implemented a more efficient queue-based parallel processing model for pystripe and added PNG reading and damaged-image-detection-and-replacement functionality. Queue-based parallel processing model in contrast to mapfunction-based is more efficient since there is no need to spawn a new worker for each image. Instead, each worker runs only one time and processes a queue of images. The queue-based model is specifically more beneficial on Windows, which has more overhead to spawn workers compared with Linux which forks them most of the time.

Using Numba[27] and NumExpr [17], we optimized the pipeline by the just-in-time compilation of some of the functions where beneficial. Also, we implemented a fast blank image detection algorithm to skip the processing of such images to further speed up the process.

**Supplementary Note 2** *Parastitcher:* We implemented a multi-GPU accelerated and NAS-friendly version of Parastitcher[11] on Linux to find the alignment of tiles. We patched TeraStitcher to repeat the read request up to 40 times to fix an issue with some NAS storage in which the reading request fails sometimes.

**Supplementary Note 3** *TSV:* We also employed a queuebased parallel processing model for the TeraStitcher volume program[44] that is more resilient to missing tiles. We increased the resiliency of TSV in case of missing tiles; TSV can now detect missing tiles and replace them with blank images. We also made TSV compatible with newer versions of Python (v3.11 at the time of publication).

**Supplementary Note 4** *16-bit to 8-bit conversion with variable bit shifting:* lightsheet fluorescent data has an unsigned 16-bit int type and brightness values have a gamma distribution. Traditional 8-bit conversion will shift the bits to the right for 8 bits, which assigns any value between 0 to 255 to zero while over-representing the tail of the gamma distribution. It is possible to clip the tail of the data so that the 16-bit image can be shifted to the right with a smaller number of bits, which leads to the assignment of a narrower range to zero. For example, with 1 right bitshift only 0 to 1 will be assigned to zero in 8-bit. We used a multi-class Otsu thresholding function from the scikit-image package[45] with 4 classes that are applied to the log1p transform of the image. Then we found 99.9 percentile of values larger than the upper threshold to find the upper bound clipping value. For the lower bound of the data, we made sure any value larger than one that could be set to zero by bit-shifting was assigned to 1 instead.

**Supplementary Note 4** *Non-blocking Paraconverter:* We used Paraconverter to convert stitched images to TeraFly format[12]. Since Paraconverter did not scale well on our cluster computer, we implemented a parallel processing algorithm that runs Paraconverter without blocking the progress of stitching for the remaining channels. We used TeraFly files for seed detection, visualization of the images in virtual reality (TeraVR)[47], and vetting of neuronal reconstruction.

**Supplementary Note 5** *Multi-channel images:* we developed an algorithm that finds the translation of the stitched images for a reference channel (usually the nuclear staining channel that is used for image registration to 3D brain atlases) at the middle z-frame of the volumes using the OpenCV version of enhanced correlation coefficient maximization algorithm[19] before converting the images to RGB format. Finally, the 2D RGB tif series are converted to a single Imaris volume with a custom code in Python.

**Supplementary Note 6** *Automation:* We minimized human interaction by automatically calculating the number of processes based on the required RAM and the available resources. Also, we made sure the RAM usage was optimal by avoiding declaring unnecessary temporary variables since the size of each z-step of the stitched images can be in the order of several GBs in RAM.

**Supplementary Note 7** *DeBleach:* `coif15` wavelet is used for initial bidirectional destriping. We used sosfiltfilt function from SciPy Signal package[38] with a frequency of (1/tile size) for generating the low-pass filter because it has no phase shift. The foreground of the image was clipped to the max of the background in advance.

**Supplementary Note 8** *Deconvolution:* We optimized the deconvolution algorithm to minimize copying data back and forth between system RAM and video RAM (vRAM), which enhanced the performance significantly. We also optimized block size calculation to ensure the maximum block size on vRAM so that larger chunks of the image volume can be processed at any given time. Image blocks now load asynchronously in the background to maximize GPU utilization time.

For an unknown reason, some of the lightsheet images could show a duplication effect on the z-axis, or neurites could look like halo tubes, which cause problems for reconstruction. Empirically, we found that a 3D Gaussian filter with sigmas of [0.5, 0.5, 1.0] and a kernel size of [3, 3, 9] on x, y, and z axes respectively, can mitigate the issue post hoc. Deconvolution algorithms are sensitive to noise. 3D Gaussian filter can also further denoise the images to improve deconvolution.

## Appendix Artifact Description

**Artifact Description (AD)**

### I. OVERVIEW OF CONTRIBUTIONS AND ARTIFACTS

#### A. Paper’s Main Contributions

*C*_1_ We introduce several novel 3D and adaptive methods and accomplish enhancement, artifact removal, segmentation and reconstruction of high-scale images.
*C*_2_ We achieve state-of-the-art (SOTA) performance in space and time even when compared to the equivalent 2D SOTA via linear complexity algorithms, which is critical for rhizomatic structures (neurons) that are power-law in size.
*C*_3_ We advance spatially sparse methodologies to enable the long-range contexts inherent in neuroscience to be computed entirely in-memory on a single workstation.
*C*_4_ Across many of our workloads, we have the first hardware-accelerated and parallel implementations for multi-GPU or multi-core environments.
*C*_5_ We enable efficient scaling to whole brain volumes (13000×13000×16000 voxels), a 67× end-to-end size increase over comparative algorithms.

#### B. Computational Artifacts

*A*_1_ https://github.com/ucla-brain/image-preprocessingpipeline
*A*_2_ https://github.com/UCLA-VAST/recut

| Artifact ID | Contributions Supported | Related Paper Elements |
| --- | --- | --- |
| $A_1$ | $C_1$ | Tables 6,8 |
| | $C_4$ | Figure 2 |
| | $C_5$ | Figure 1 |
| $A_2$ | $C_1$ | Tables 1-7 |
| | $C_2$ | Figure 3,4 |
| | $C_3$ | Figure 4 |
| | $C_4$ | Figure 3 |
| | $C_5$ | Figure 1 |

### II. ARTIFACT IDENTIFICATION

#### A. Computational Artifact A_1_

##### Relation To Contributions

*A*_1_ is the software for preprocessing images. All images are preprocessed with this software therefore all accuracy and even runtime results are affected by this preparation procedure. Table 8 column (raw image unprocessed) however, indicates the accuracy *without* using the main aspects of the preprocessing procedure (still uses stitching).

##### Expected Results

Images preprocessed with *A*_1_ have qualitatively lower errors as shown in Figure 1 and the signal to noise ratio is higher which can be noted quantitatively, though indirectly, by the compression ratios in Figure 2. *A*_1_ can run on the two provided datasets *D*_1_ and *D*_2_, alternate methods or implementations simply crash at these image sizes due to scale. Due to the multi-GPU and multi-CPU implementation, *A*_1_ is SOTA in runtime and throughput.

##### Expected Reproduction Time (in Minutes)

All runtimes are recorded on the recommended system listed below. It takes 2 days per brain to conduct destriping, stitching and debleaching. Finally, it takes 360 minutes to deconvolve each dataset.

##### Artifact Setup (incl. Inputs)

*Hardware:* For these workloads, we recommend a single workstation with 2× AMD EPYC 7763 (128 cores/ 256 threads) CPUs, 4 TB RAM. and 8× A100 GPUs, connected with NVSwitch. The large memory capacity allows data sizes of 1 whole mouse brain at 1um^3^ resolution to be processed entirely in-memory for all tasks.

##### Software

The software page https://github.com/uclabrain/image-preprocessing-pipeline renders the README.md which gives explicit instructions on software packages and installation.

##### Datasets / Inputs

*D*_1_ is whole brain image at .4 um^3^ resolution (6x obj. lens). We provide only the fully preprocessed version of the brain to reproduce the results. *D*_2_ is a half brain (hemisphere) at 1 um^3^ resolution (15x). *D*_2_ is available in raw and preprocessed forms. We are in the process of making both datasets publicly available through the brain image library a nationally funded cloud specialized in storing terabyte scale brain volumes such as those used in this paper.

##### Installation and Deployment

The software page https://github.com/ucla-brain/image-preprocessing-pipeline renders the README.md which gives explicit instructions on installation and usage. Usage details show how to transform *D*_2_ from unprocessed to preprocessed to reproduce the results in the paper related to image artifacts.

##### Artifact Execution

The workflow and tasks of preprocessing are laid out explicitly with additional figures and inputs and outputs on the same web page: https://github.com/ucla-brain/image-preprocessing-pipeline. Please use the default parameters and settings. All timings reported in the paper are from a single run due to their adequate stability and long runtime.

##### Artifact Analysis (incl. Outputs)

*A*_1_ takes images in various formats and outputs images with known artifacts removed. To see the full list of supported inputs and outputs refer to Figure 1 in the paper.

#### B. Computational Artifact A_2_

##### Relation To Contributions

*A*_2_ is the software for conversion of images to VDB, semantic segmentation and skeletonization. All major contributions of the paper are either analyzed, mediated or produced by this software. Note, this software also provides TreeBench the software for evaluating the accuracy Tables 6-8 though Table 8 only relates to ablations in preprocessing. In contrast to *A*_1_, this software is only dual CPU in a single node, it does not use GPU.

##### Expected Results

*A*_2_ (Recut) is instrumented to report the runtimes of its individual stages (see Figure 1), which are then collected from logs and plotted in the scripts. Recut essentially transforms images into skeletons of neurons. These neurons are then compared to their ground truth (corrected) counterpart to analyze accuracy.

##### Expected Reproduction Time (in Minutes)

On our example system, conversion and skeletonization combined takes 66 minutes. Of this time, conversion takes 21 minutes, seed segmentation takes 35 minutes, cell segmentation takes 9 minutes and skeletonization takes 2 minutes.

##### Artifact Setup (incl. Inputs)

###### Hardware

For these workloads, we recommend a single workstation with 2× AMD EPYC 7763 (128 cores/ 256 threads) CPUs and 4 TB RAM. 4TB keeps workloads in-memory, despite large peaks in dynamic memory with the (optional) 256-threads in flight. Systems with less cores inherently require less dynamic memory. However, one can always lower the maximum thread count via command line parameters to reduce the required system memory further.

###### Software, Installation and Deployment

Installing Recut involves 2 commands, installing a package manager and installing the recut package. These are listed on the software page https://github.com/UCLA-VAST/recut. You do not need to concern yourself with languages or packages, this works on Windows Subsystem for Linux and any native Linux distribution but has not been tested on MacOS. We use the Nix package manager since it is the gold standard in reproducible and portable builds. It is far more strict in terms of build determinism than Docker or other similar technologies.

###### Datasets / Inputs

Image datasets are the same as above. However, *A*_2_ also refers to a set of ground truth skeletons for each image. We also extract the cell bodies from ground truth skeletons as inputs to skeletonization runs to isolate accuracy related to branch reconstruction (see the third command below).

###### Artifact Execution

We can split up the tasks of reconstruction into conversion, seed segmentation and skeletonization when benchmarking accuracy and performance. Running these steps is explained more in the Github Readme as well but for easy reference run:

~~~
recut [tif_folder] --output-type mask
     --voxel-size .4 .4 .4
recut [mask vdb from previous conversion step]
     --output-type seeds --voxel-size .4 .4 .4 recut [same mask vdb] --seeds
[ground truth skeleton folder]
     --voxel-size .4 .4 .4
~~~

###### Artifact Analysis (incl. Outputs)

The outputs of reconstruction are namely compressed images (VDBs), seeds or skeletons. If you do not have a system that can run these commands for these image sizes, we can provide all of the logs and skeletons from the runs used in the paper.

After a reconstruction of skeletons runs you will notice a new folder in the directory named run-1. This folder has a set of subfolders for each connected component. Some connected components contain multiple seeds (folders). In the process of skeletonization, graphs are partitioned into trees rooted at seeds, therefore components with multiple seeds will have multiple skeletons (.swc) files.

We have python scripts in the repositories scripts folder which explain how to install DIADEM, the 1 external dependency of analysis. These scripts produce the accuracy tables 5-8, and the sequential runtime plot and memory usage plot (Figure 3-4).

## References

[1] Ludovica Acciai, Paolo Soda, and Giulio Iannello. 2016. Automated Neuron Tracing Methods: An Updated Account. Neuroinformatics (2016). 10.1007/s12021-016-9310-0

[2] Nanda S. Maraver P. et al. Akram, M. 2018. An open repository for single-cell reconstructions of the brain forest. Sci. Data 5, Article 18006 (2018). 10.1038/sdata.2018.6

[3] Katrin Amunts and Thomas Lippert. 2021. Brain research challenges supercomputing. Science 374, 6571 (2021), 1054–1055.abl8519 10.1126/science.

[4] Cameron Arshadi, Ulrik Günther, Mark Eddison, Kyle I. S. Harrington, and Tiago A. Ferreira. 2021. SNT: a unifying toolbox for quantification of neuronal anatomy. Nature Methods 18, 4 (apr 2021), 374–377. 10.1038/s41592-021-01105-7

[5] Giorgio A. Ascoli, Duncan E. Donohue, and Maryam Halavi. 2007. NeuroMorpho.Org: A Central Resource for Neuronal Morphologies. Journal of Neuroscience 27, 35 (2007), 9247–9251. 10.1523/JNEUROSCI.2055-07.2007 arXiv:https://www.jneurosci.org/content/27/35/9247.full.pdf

[6] Andreas Bærentzen, Rasmus Emil Christensen, Emil Toftegaard Gæde, and Eva Rotenberg. 2023. Multilevel Skeletonization Using Local Separators. arXiv preprint 2303.07210 (2023).

[7] Andreas Bærentzen and Eva Rotenberg. 2021. Skeletonization via Local Separators. ACM Trans. Graph. 40, 5, Article 187 (sep 2021), 18 pages. 10.1145/3459233

[8] R. Bayer and E.M. McCreight. 1972. Organization and maintenance of large ordered indexes. Acta Informatica 1 (1972), 173–189. 10.1007/BF00288683

[9] Klaus Becker, Saiedeh Saghafi, Marko Pende, Inna Sabdyusheva-Litschauer, Christian M Hahn, Massih Foroughipour, Nina Jährling, and Hans-Ulrich Dodt. 2019. Deconvolution of light sheet microscopy recordings. Scientific reports 9, 1 (2019), 17625.

[10] George Bezerra, Divy Agrawal, and Amr El Abbadi. 2013. X-Stream: Edge-centric Graph Processing using Streaming Partitions. in Proceedings of the 24th ACM Symposium on Operating Systems Principles (SOSP). ACM, 472–488. 10.1145/2517349.2522740

[11] Alessandro Bria and Giulio Iannello. 2012. TeraStitchera tool for fast automatic 3D-stitching of teravoxel-sized microscopy images. BMC bioinformatics 13, 1 (2012), 1–15.

[12] Alessandro Bria, Giulio Iannello, Leonardo Onofri, and Hanchuan Peng. 2016. TeraFly: real-time three-dimensional visualization and annotation of terabytes of multidimensional volumetric images. Nature methods 13, 3 (2016), 192–194.

[13] Alessandro Bria, Giulio Iannello, Leonardo Onofri, and Hanchuan Peng. 2022. TeraFly: real-time three-dimensional visualization and annotation of terabytes of multidimensional volumetric images. 13 (2022). Issue 3. 10.1038/nmeth.3767

[14] Robert C. Cannon, Dennis A. Turner, G.K. Pyapali, and H.V. Wheal. 1998. An on-line archive of reconstructed hippocampal neurons. Journal of Neuroscience Methods 84, 1-2 (1998), 49–54. 10.1016/S0165-0270(98)00091-0

[15] Angel X. Chang, Thomas A. Funkhouser, Leonidas J. Guibas, Pat Hanrahan, Qi-Xing Huang, Zimo Li, Silvio Savarese, Manolis Savva, Shuran Song, Hao Su, Jianxiong Xiao, Li Yi, and Fisher Yu. 2015. ShapeNet: An Information-Rich 3D Model Repository. CoRR abs/1512.03012 (2015). 1512.03012 http://arxiv.org/abs/1512.03012

[16] Yuze Chi, Guohao Dai, Yu Wang, Guangyu Sun, Guoliang Li, and Huazhong Yang. 2015. NXgraph: An Efficient Graph Processing System on a Single Machine. CoRR abs/1510.06916 (2015). 1510.06916 http://arxiv.org/abs/1510.06916

[17] D Cooke, T Hochberg, F Alted, I Vilata, M Wiebe, G de Menten, A Valentino, and RA McLeod. 2009. NumExpr: Fast numerical expression evaluator for NumPy.

[18] Nicu D Cornea, Deborah Silver, and Patrick Min. 2005. Curve-skeleton applications. in VIS 05. IEEE Visualization, 2005. IEEE, 95–102.

[19] Georgios D Evangelidis and Emmanouil Z Psarakis. 2008. Parametric image alignment using enhanced correlation coefficient maximization. IEEE transactions on pattern analysis and machine intelligence 30, 10 (2008), 1858–1865.

[20] Linqing Feng, Ting Zhao, and Jinhyun Kim. 2015. neu-Tube 1.0: A New Design for Efficient Neuron Recon-struction Software Based on the SWC Format. eNeuro 2, 1 (January 2015), ENEURO.0049–14.2014. 10.1523/eneuro.0049-14.2014

[21] Le Gao, Sang Liu, Lingfeng Gou, and Yachuang Hu et al. 2022. Single-neuron projectome of mouse prefrontal cortex. 25 (2022), 515–529. Issue 4. 10.1038/s41593-022-01041-5

[22] T.A. Gillette, K.M. Brown, and G.A. Ascoli. 2011. The DIADEM Metric: Comparing Multiple Reconstructions of the Same Neuron. Neuroinformatics 9 (2011), 233–245. 10.1007/s12021-011-9117-y

[23] Patric Hellermann. 2023. Neural radiance fields and billion-dollar opportunities they create in the 3D tool stack. https://foundamental.com/aecvc/nerfs-in-3d

[24] Yuanming Hu, Tzu-Mao Li, Luke Anderson, Jonathan Ragan-Kelley, and Frédo Durand. 2019. Taichi: a language for high-performance computation on spatially sparse data structures. ACM Transactions on Graphics (TOG) 38, 6 (2019), 201.

[25] Shengdian Jiang, Yimin Wang, Lijuan Liu, Sujun Zhao, Mengya Chen, Xuan Zhao, Peng Xie, Liya Ding, Zongcai Ruan, Hong-Wei Dong, Giorgio A. Ascoli, Michael Hawrylycz, Hongkui Zeng, and Hanchuan Peng. 2021. MorphoHub: A Platform for Petabyte-Scale Multi-Morphometry Generation. bioRxiv (2021). 10.1101/2021.01.09.426010

[26] Aapo Kyrola, Guy Blelloch, and Carlos Guestrin. 2012. GraphChi: Large-Scale Graph Computation on Just a PC. in Proceedings of the 10th USENIX Conference on Operating Systems Design and Implementation (OSDI’12). USENIX Association, Berkeley, CA, USA, 31–46. https://www.usenix.org/conference/osdi12/technical-sessions/presentation/kyrola

[27] Siu Kwan Lam, Antoine Pitrou, and Stanley Seibert. 2015. Numba: A llvm-based python jit compiler. in Proceedings of the Second Workshop on the LLVM Compiler Infrastructure in HPC. 1–6.

[28] Hsueh-Ti Derek Liu, Francis Williams, Alec Jacobson, Sanja Fidler, and Or Litany. 2022. Learning Smooth Neural Functions via Lipschitz Regularization. 10.48550/ARXIV.2202.08345

[29] Yufeng Liu, Shengdian Jiang, Yingxin Li, Sujun Zhao, Zhixi Yun, Lijuan Liu, Hanchuan Peng, et al. 2023. Full-Spectrum Neuronal Diversity and Stereotypy through Whole Brain Morphometry. Research Square (2023).

[30] Chiara Magliaro, Alejandro L. Callara, Nicola Vanello, and Arti Ahluwalia. 2019. Gotta trace ‘em all: A mini-review on tools and procedures for segmenting single neurons toward deciphering the structural connectome. Frontiers in Bioengineering and Biotechnology 7, AUG (2019), 1–8. 10.3389/fbioe.2019.00202

[31] Linus Manubens-Gil, Zhi Zhou, Hanbo Chen, Arvind Ramanathan, Xiaoxiao Liu, Yufeng Liu, Alessandro Bria, Todd Gillette, Zongcai Ruan, Jian Yang, Miroslav Radojević, Ting Zhao, Li Cheng, Lei Qu, Siqi Liu, Kristofer E. Bouchard, Lin Gu, Weidong Cai, Shuiwang Ji, Badrinath Roysam, Ching-Wei Wang, Hongchuan Yu, Amos Sironi, Daniel Maxim Iascone, Jie Zhou, Erhan Bas, Eduardo Conde-Sousa, Paulo Aguiar, Xiang Li, Yujie Li, Sumit Nanda, Yuan Wang, Leila Muresan, Pascal Fua, Bing Ye, Hai yan He, Jochen F. Staiger, Manuel Peter, Daniel N. Cox, Michel Simonneau, Marcel Oberlaender, Gregory Jefferis, Kei Ito, Paloma Gonzalez-Bellido, Jinhyun Kim, Edwin Rubel, Hollis T. Cline, Hongkui Zeng, Aljoscha Nern, Ann-Shyn Chiang, Jianhua Yao, Jane Roskams, Rick Livesey, Janine Stevens, Tianming Liu, Chinh Dang, Yike Guo, Ning Zhong, Georgia Tourassi, Sean Hill, Michael Hawrylycz, Christof Koch, Erik Meijering, Giorgio A. Ascoli, and Hanchuan Peng. 2022. BigNeuron: A resource to benchmark and predict best-performing algorithms for automated reconstruction of neuronal morphology. bioRxiv (2022). 10.1101/2022.05.10.491406

[32] Karl Marrett, Muye Zhu, Yuze Chi, Chris Choi, Zhe Chen, Hong-Wei Dong, Chang Sin Park, X. William Yang, and Jason Cong. 2021. Recut: a Concurrent Framework for Sparse Reconstruction of Neuronal Morphology. bioRxiv (2021). 10.1101/2021.12.07.471686

[33] Gokhan Memik, Dipanjan Sengupta, Emre Kultursay, Nandhini Chandramoorthy, and Gokhan Memik. 2013. MapGraph: A High Level Framework for Graph Processing on GPUs. Proceedings of the 18th International Conference on Architectural Support for Programming Languages and Operating Systems (ASPLOS ‘13) (2013), 153–166. 10.1145/2451116.2451132

[34] Thomas Müller, Alex Evans, Christoph Schied, and Alexander Keller. 2022. Instant Neural Graphics Primitives with a Multiresolution Hash Encoding. ACM Trans. Graph. 41, 4, Article 102 (2022), 15 pages. 10.1145/3528223.3530127

[35] Ken Museth. 2013. VDB: High-Resolution Sparse Volumes with Dynamic Topology. ACM Trans. Graph. 32, 3, Article 27 (2013), 22 pages. 10.1145/2487228.2487235

[36] Ken Museth. 2017. Novel Algorithm for Sparse and Parallel Fast Sweeping: Efficient Computation of Sparse Signed Distance Fields. in ACM SIGGRAPH 2017 Talks (Los Angeles, California) (SIGGRAPH ‘17). Association for Computing Machinery, New York, NY, USA, Article 74, 2 pages. 10.1145/3084363.3085093

[37] Lawrence Page, Sergey Brin, Rajeev Motwani, and Terry Winograd. 1998. The PageRank Citation Ranking: Bringing Order to the Web. Technical Report. Stanford Digital Library Technologies Project. http://citeseerx.ist.psu.edu/viewdoc/summary?doi=10.1.1.31.1768

[38] Ashwin Pajankar. 2017. Signal processing with SciPy. Raspberry Pi Supercomputing and Scientific Programming: MPI4PY, NumPy, and SciPy for Enthusiasts (2017), 139–147.

[39] Pietro Ricci, Vladislav Gavryusev, Caroline Müllenbroich, Lapo Turrini, Giuseppe de Vito, Ludovico Silvestri, Giuseppe Sancataldo, and Francesco Saverio Pavone. 2022. Removing striping artifacts in lightsheet fluorescence microscopy: a review. Progress in Biophysics and Molecular Biology 168 (2022), 52–65. 10.1016/j.pbiomolbio.2021.07.003 The Resolution Revolution: Fluorescence Microscopy of Biological Samples from Micro to Meso.

[40] Olaf Ronneberger, Philipp Fischer, and Thomas Brox. 2015. U-Net: Convolutional Networks for Biomedical Image Segmentation. CoRR abs/1505.04597 (2015). 1505.04597 http://arxiv.org/abs/1505.04597

[41] Darius Rückert, Yuanhao Wang, Rui Li, Ramzi Idoughi, and Wolfgang Heidrich. 2022. NeAT: Neural Adaptive Tomography. ACM Trans. Graph. 41, 4, Article 55 (2022), 13 pages. 10.1145/3528223.3530121

[42] James A. Sethian. 1996. A fast marching level set method for monotonically advancing fronts. Proc. of the National Academy of Sciences of the USA (Feb 1996), 1591–1595.

[43] Ernst HK Stelzer, Frederic Strobl, Bo-Jui Chang, Friedrich Preusser, Stephan Preibisch, Katie McDole, and Reto Fiolka. 2021. Light sheet fluorescence microscopy. Nature Reviews Methods Primers 1, 1 (2021), 73.

[44] Justin Swaney, Lee Kamentsky, Nicholas B Evans, Katherine Xie, Young-Gyun Park, Gabrielle Drummond, Dae Hee Yun, and Kwanghun Chung. 2019. Scalable image processing techniques for quantitative analysis of volumetric biological images from light-sheet microscopy. BioRxiv (2019), 576595.

[45] Stefan Van der Walt, Johannes L Schönberger, Juan Nunez-Iglesias, François Boulogne, Joshua D Warner, Neil Yager, Emmanuelle Gouillart, and Tony Yu. 2014. scikit-image: image processing in Python. PeerJ 2 (2014), e453.

[46] Matthew B. Veldman, Chang Sin Park, Charles M. Eyermann, Jason Y. Zhang, Elizabeth Zuniga-Sanchez, Arlene A. Hirano, Tanya L. Daigle, Nicholas N. Foster, Muye Zhu, Peter Langfelder, Ivan A. Lopez, Nicholas C. Brecha, S. Lawrence Zipursky, Hongkui Zeng, Hong-Wei Dong, and X. William Yang. 2020. Brainwide Genetic Sparse Cell Labeling to Illuminate the Morphology of Neurons and Glia with Cre-Dependent MORF Mice. Neuron 108, 1 (2020), 111–127.e6. 10.1016/j.neuron.2020.07.019

[47] Yimin Wang, Qi Li, Lijuan Liu, Zhi Zhou, Zongcai Ruan, Lingsheng Kong, Yaoyao Li, Yun Wang, Ning Zhong, Renjie Chai, et al. 2019. TeraVR empowers precise reconstruction of complete 3-D neuronal morphology in the whole brain. Nature communications 10, 1 (2019), 3474.

[48] Johan Winnubst, Erhan Bas, Tiago A. Ferreira, Nelson Spruston, Karel Svoboda, and Jayaram Chandrashekar. 2019. Reconstruction of 1,000 Projection Neurons Reveals New Cell Types and Organization of Long-Range Connectivity in the Mouse Brain. Cell 179, 1 (2019), 268–281.e13. 10.1016/j.cell.2019.07.042

[49] Hang Xiao and Hanchuan Peng. 2013. APP2: Automatic tracing of 3D neuron morphology based on hierarchical pruning of a gray-weighted image distance-tree. Bioinformatics 29, 11 (2013), 1448–1454. 10.1093/bioinformatics/btt170

[50] Jian Yang, Ming Hao, Xiaoyang Liu, Zhijiang Wan, Ning Zhong, and Hanchuan Peng. 2019. FMST: an Automatic Neuron Tracing Method Based on Fast Marching and Minimum Spanning Tree. Neuroinformatics 17, 2 (2019), 185–196. 10.1007/s12021-018-9392-y

[51] Hongkui Zeng and Joshua R Sanes. 2017. Neuronal cell-type classification: challenges, opportunities and the path forward. Nature Publishing Group 18 (2017). 10.1038/nrn.2017.85

[52] Hongkai Zhao. 2004. Fast Sweeping Method for Eikonal Equations. Math. Comp. 74 (2004), 603–627.

[53] Hongkai Zhao. 2007. Parallel implementation of the fast sweeping method. Journal of Computational Mathematics 25 (2007), 421–429. Issue 4.

